# P2Y6 receptor signaling in natural killer cells impairs insulin sensitivity in obesity

**DOI:** 10.1101/2021.06.23.449596

**Authors:** Anna Sieben, Anne Christine Hausen, Paul Klemm, Alexander Jais, Christian Heilinger, Jens Alber, Kat Folz-Donahue, Lena Schumacher, Janine Altmüller, Marek Franitza, Patrick Giavalisco, Yvonne Hinze, Susanne Brodesser, Christian Kukat, Sebastian Theurich, Jens C. Brüning

**Author notes:** Address correspondence to: Jens C. Brüning, M.D., Max Planck Institute for Metabolism Research, Gleueler Str. 50, 50931 Cologne, Germany.

## Abstract

Natural killer (NK) cells contribute to the development of obesity-associated insulin resistance and have previously been shown to up-regulate the expression of the P2Y purinoreceptor 6 (P2Y6R) upon high fat diet (HFD)-induced obesity. Here, we reveal that NK cell-specific inactivation of the P2Y6R gene improves insulin sensitivity in obese mice and reduces the expression of chemokines in adipose tissue infiltrating NK-cells. Obese mice lacking P2Y6R specifically in NK cells exhibited a reduction in adipose tissue inflammation, exhibited improved insulin-stimulated suppression of lipolysis in adipose tissue and a reduction in hepatic glucose production, leading to an overall improvement of systemic insulin sensitivity. In contrast, myeloid lineage specific P2Y6R inactivation does not affect energy or glucose homeostasis in obesity. Collectively, we show that P2Y6R signaling in NK cells contributes to the development of obesity-associated insulin resistance and thus might be a future target for the treatment of obesity-associated insulin resistance and type 2 diabetes.

## Introduction

More than 1.9 billion adults were overweight in 2016. Of these, over 650 million adults were obese (WHO, 2021), reflecting a slow but dangerous global pandemic that started more than 40 years ago (WHO, 2021). Obesity not only increases the risk of co-morbidities such as cardiovascular diseases, cancer and diabetes mellitus (Guh *et al*., 2009) but also has profound effects on the immune system (De Heredia *et al*., 2012). Obesity is associated with a chronic low-grade inflammation, termed metaflammation (De Heredia *et al*., 2012), which is reflected in an increased number of macrophages (Weisberg *et al*., 2003), T cells and natural killer (NK) cells (Theurich *et al*., 2017) infiltrating the adipose tissue and a rise in pro-inflammatory molecules in tissues and blood (Xu *et al*., 2003; Schmidt *et al*., 2015).

Nucleotides such as adenosine and uridine are critical building blocks of the genetic code and at the same time key regulators of inflammation (Idzko *et al*., 2014). Uridine acts as a potent anti-inflammatory molecule during lung inflammation (Evaldsson *et al*., 2007) and pulmonary fibrosis (Cicko *et al*., 2015). Here, it affects adhesion between leukocytes and endothelium in a lung inflammation model (Uppugunduri and Gautam, 2004). Plasma uridine concentrations are regulated by fasting and refeeding (Deng *et al*., 2017), controlled by the circadian rhythm (El Kouni *et al*., 1990) and exercise (Yamamoto *et al*., 1997) and positively correlate with insulin resistance and plasma insulin levels (Hamada *et al*., 2007). Uridine impairs insulin sensitivity and glucose tolerance in mice (Urasaki *et al*., 2014), has an impact on liver lipid metabolism (Urasaki *et al*., 2016; Le *et al*., 2013) as well as is involved in glycogen metabolism (Haugaard *et al*., 1977).

Moreover, uridine serves as precursor for the synthesis of the nucleotide uridine diphosphate (UDP). Coupled to glucose, this nucleotide is a critical component in the synthesis pathway of glycogen (Berg *et al*., 2002). At the same time, UDP acts as an extracellular signaling molecule after being released from apoptotic and necrotic cells and serves as danger and find-me signal in order to attract immune cells to the site of inflammation (Elliott *et al*., 2009). On the other hand, a controlled way of nucleotide release has been reported (Lazarowski *et al*., 2011). Here, the extracellular nucleotide adenosine triphosphate (ATP) can act as an initiator or terminator of immune responses to induce different effects on immune cells depending on the pattern of P2 receptors engaged, the duration of the stimulus and its concentration in the extracellular milieu (Gorini *et al*., 2010).

The extracellular nucleotide UDP binds to P2Y6 receptors (P2Y6R) which belong to the family of G-protein coupled receptors. The activation of Gq-coupled P2Y6Rs results in a rise of intracellular calcium concentration leading to an altered activity of kinases such as protein kinase C (PKC), mitogen-activated protein kinase (MAPK) and extracellular signal-regulated kinase (ERK), to changes in transcription and to the release of signaling molecules (Kim *et al*., 2003; Li *et al*., 2014; Zhang *et al*., 2011). In the context of glucose metabolism and obesity it has been shown that UDP-P2Y6R signaling regulates feeding via hypothalamic agouti-related protein (AgRP) neurons (Steculorum *et al*., 2015) and that P2Y6R action in AgRP neurons contributes to HFD-induced deterioration of insulin sensitivity (Steculorum *et al*., 2017). Investigations in peripheral organs revealed that adipocyte-specific deletion of P2Y6R protects mice from diet-induced obesity, improves glucose tolerance and insulin sensitivity and reduces systemic inflammation (Jain *et al*., 2020). In contrast to that, P2Y6R deletion in skeletal muscle resulted in impaired glucose homeostasis (Jain *et al*., 2020).

P2Y6Rs are also expressed in a variety of immune cells, such as macrophages (Bar *et al*., 2008; Lattin *et al*., 2008), T cells (Giannattasio *et al*., 2011) and microglia (Idzko *et al.,* 2004, Kim *et al.,* 2011). Moreover, expression of P2Y6R is upregulated in NK cells isolated from adipose tissue of mice fed a HFD when compared to lean mice on a control diet (Theurich *et al*., 2017). NK cells are part of the innate immune system responsible for the recognition and elimination of virus-infected and cancerous cells (Vivier *et al*., 2008). They are able to destroy target cells via the release and joined action of the cytotoxic mediators perforin and granzyme (Pardo *et al*., 2002) or through the exocytosis of cytokines such as interferon gamma (IFNγ) and tumor necrosis factor alpha (TNFα) (Girart *et al*., 2007; Wang *et al*., 2012; Jewett *et al*., 1996). NK cell activity is controlled by the orchestration of activating and inactivating receptors (Bakker *et al*., 2000; Lanier, 2008) as well as by cytokines such as interleukin (IL) 2 and 15 (Zwirner and Domaica, 2010). Obesity induces lipid accumulation in NK cells, leading to cytotoxic dysfunction, impaired trafficking and reduced anti-tumor responses (Michelet *et al*., 2018). NK cells derived from adipose tissue of obese mice show an activated immune phenotype and release cytokines that trigger the differentiation and infiltration of macrophages, finally resulting in increased inflammation and promotion of insulin resistance (Wensveen *et al*., 2015; Lee *et al*., 2016). The majority of macrophages infiltrating the adipose tissue of obese animals and humans are arranged around dead adipocytes, forming characteristic crown-like structures (CLS) (Murano *et al*., 2008; Cinti *et al*., 2005). These necrotic adipocytes might release UDP as danger signal leading to an increased infiltration or local proliferation of immune cells, thus promoting metaflammation and development of insulin resistance.

Given the improvement of insulin action in conventional P2Y6R knockout mice (Steculorum *et al*., 2017; Jain *et al*., 2020), and the role of P2Y6R-signaling in immune cells, we aimed to investigate the role of this receptor in NK cells and macrophages in the development of obesity-associated metaflammation and insulin resistance. To this end, we generated mice lacking P2Y6R specifically in these immune compartments. While ablation of P2Y6R in myeloid lineage cells had no effect on metaflammation and obesity-associated insulin resistance, obese mice lacking P2Y6R specifically in NK cells exhibited a reduction in adipose tissue inflammation, exhibited improved insulin-stimulated suppression of lipolysis in adipose tissue and a reduction in hepatic glucose production, leading to an overall improvement of systemic insulin sensitivity. Collectively, these experiments extend the tissue-specific actions of P2Y6R signaling in obesity, further characterizing P2Y6R inhibition as a novel target for metabolic diseases.

## Results

### Deletion of the P2Y6R gene in NK cells does not protect from HFD-induced obesity

Among other genes, P2Y6R expression contributes to the discrimination of NK cells from lean and obese mice (Theurich *et al*., 2017). Furthermore, it has been reported that circulating uridine levels are increased in obesity (Steculorum *et al*., 2015; Deng *et al*., 2017). To further define potential changes in uridine and UDP in different organismal compartments upon development of HFD-induced obesity and insulin resistance, we compared uridine and UDP levels in plasma, liver, adipocytes and the adipose tissue stromal vascular fraction (SVF) of mice that had been kept on normal chow diet (NCD) or had been exposed to HFD-feeding for 6 or 12 weeks. This analysis revealed increased levels of uridine in plasma of obese compared to lean mice after 12 weeks of HFD feeding (Suppl. Fig. S1A), while uridine levels in liver and SVF increased already at 6 weeks of HFD feeding and remained elevated until 12 weeks of HFD feeding (Suppl. Fig. S1B, C), while adipocyte uridine concentration increased after 6 weeks of HFD feeding, but reached comparable levels to NCD-fed mice after 12 weeks (Suppl. Fig. S1D). While UDP levels in plasma paralleled the increase of circulating uridine 12 weeks after HFD-feeding (Suppl. Fig. S1E) as well as the increases in adipose tissue SVF 6 and 12 weeks after high fat diet feeding (Suppl. Fig. S1G), hepatic UDP levels remained unaltered upon HFD feeding (Suppl. Fig. S1F), and adipocyte UDP levels increased only after prolonged HFD feeding (Suppl. Fig. S1H). These experiments indicate a coordinated regulation of uridine and UDP concentrations in plasma and SVF, while UDP synthesis in liver and adipocytes does not appear to be directly linked to changes in local uridine concentrations.

**Fig. S1:**
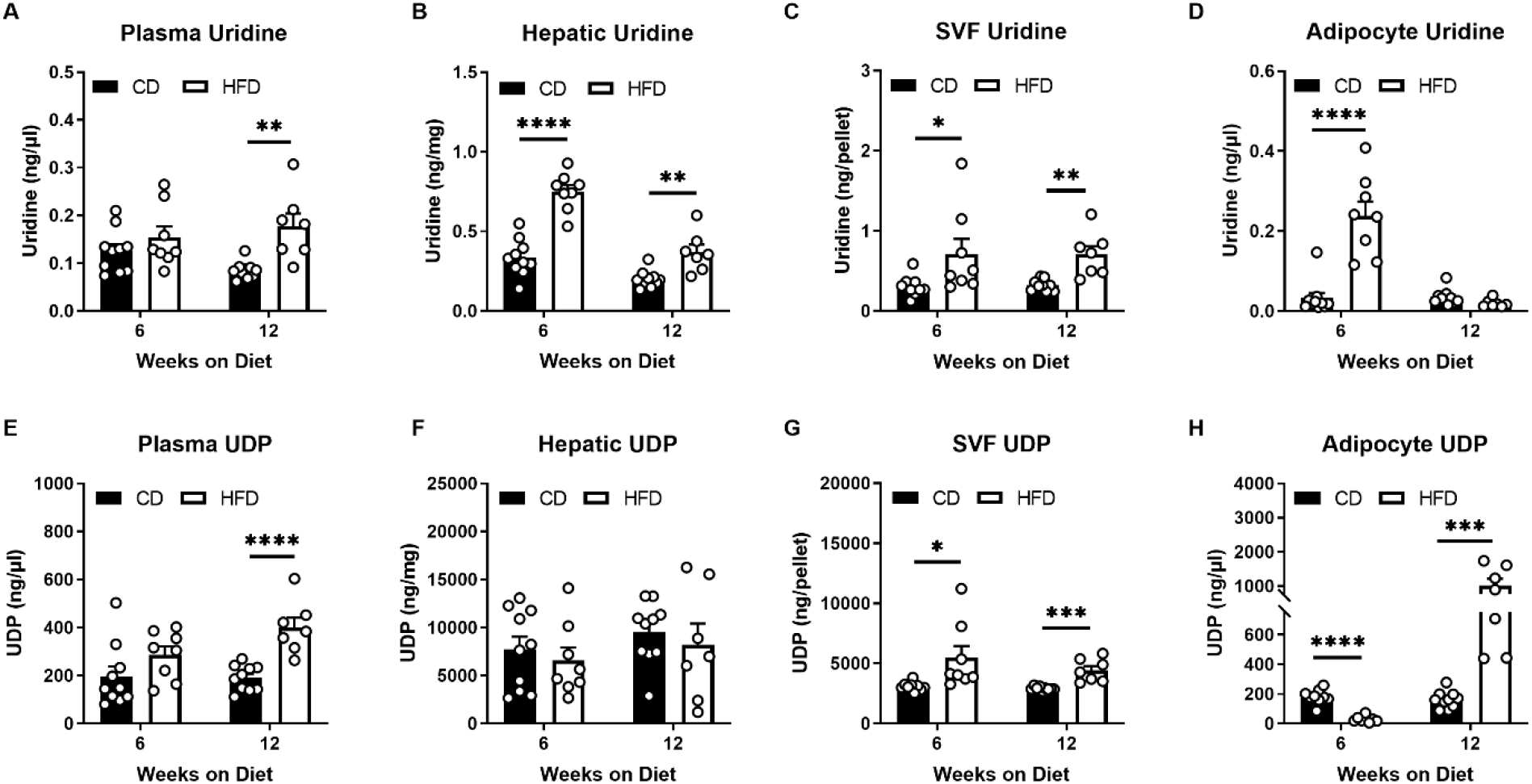
Uridine and UDP levels are increased upon HFD feeding. Two cohorts of C57Bl6 mice were either fed a normal chow diet (NCD) or HFD for 6 (NCD, n=10; HFD, n=8) or 12 (NCD, n=10; HFD, n=7) weeks. Uridine and UDP levels were determined in plasma **(A, E)**, liver **(B, F)**, cells of the stromal vascular fraction (SVF) **(C, G)** and in adipocytes **(D, H)**. Statistics: Unpaired two-tailed t-test, *p≤0.05, **p≤0.01, ***p≤0.001, ****p≤0.0001. Data are represented as mean ± SEM. ○ denotes individual mice.

In light of the coordinated regulation of uridine and UDP in plasma and adipose tissue SVF, compartments which directly interact with immune cells, we aimed to investigate the impact of P2Y6R signaling in NK cells on body weight development, glucose metabolism, and obesity-associated metaflammation *in vivo*. In order to specifically delete the P2Y6R in NK cells, we crossed mice carrying a loxP-flanked P2Y6R gene (Jain *et al*., 2020) with Ncr1-Cre mice (Eckelhardt *et al*., 2011). PCR analysis of DNA isolated from NK cells of control (P2Y6^fl/fl^) or NK cell-specific P2Y6-deleted (P2Y6^ΔNcr1^) mice confirmed successful and specific disruption of P2Y6R expression in NK cells (Suppl. Fig. S2).

**Fig. S2:**
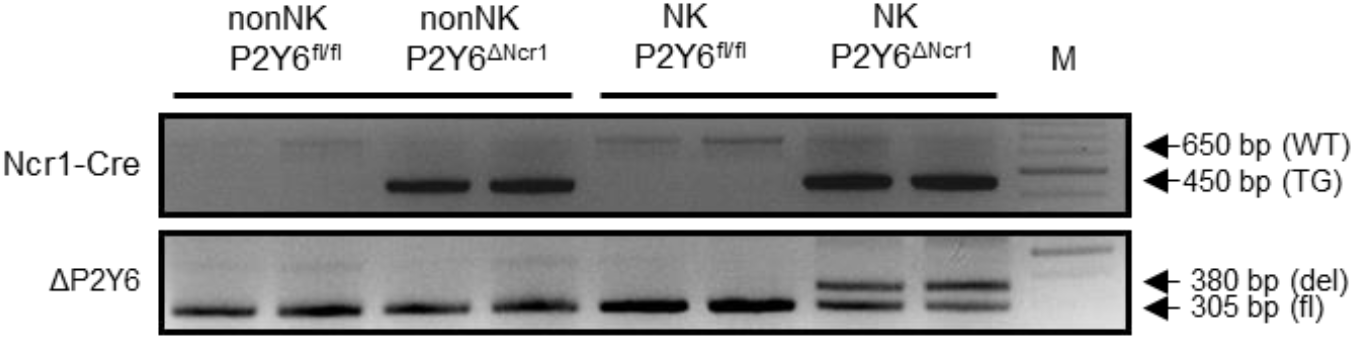
P2Y6R is selectively deleted in NK cells. Splenic cells from P2Y6^fl/fl^ (n=2) and P2Y6^ΔNcr1^ (n=2) mice were separated with magnetic beads resulting in two fractions: NK cells and nonNK cells. Genomic DNA was extracted from each group. Genotyping for P2Y6R was performed by competitive PCR using the primers listed in table 1 which resulted in a 305 bp PCR-product (floxed) and a 370 bp PCR-product (P2Y6R deletion) in NK cells isolated from P2Y6^ΔNcr1^ (Ncr1-Cre positive mice (TG band 450 bp)). This deletion band was neither detected in NK cells from P2Y6^fl/fl^, (Ncr1-Cre negative mice (WT band 650 bp)) nor in nonNK cells of either genotype.

Starting at the age of 6 weeks, P2Y6^ΔNcr1^ mice and their respective littermate controls were fed a HFD or control diet (CD). HFD feeding did not result in any differences in body weight between genotypes in male mice (Fig.1A), while female P2Y6^ΔNcr1^ mice exhibited a mild protection from HFD-induced weight gain upon prolonged HFD-feeding (Suppl. Fig. S3A). Consistent with unaltered body weight development, the relative amount of lean and fat mass of male mice after 15 weeks on HFD was not different between genotypes (Fig. 1B). In line with this, male P2Y6^ΔNcr1^ mice did not exhibit any differences in food intake (Suppl. Fig. S4A), water consumption (Suppl. Fig. S4B), energy expenditure (Fig. S4C) or locomotor activity (Fig. S4D) compared with control mice on HFD. However, after 14 weeks of HFD feeding we could observe a slight reduction in liver weights in male (Fig. 1C) and female (Suppl. Fig. S3B) P2Y6^ΔNcr1^ mice, as well as a decrease in fasted leptin levels in male P2Y6^ΔNcr1^ mice compared to their control littermates (Fig. 1D). Collectively, NK-cell P2Y6R signaling has no major impact on energy homeostasis and body weight control in obesity, while it appears to affect liver weight in obesity.

**Fig. S3:**
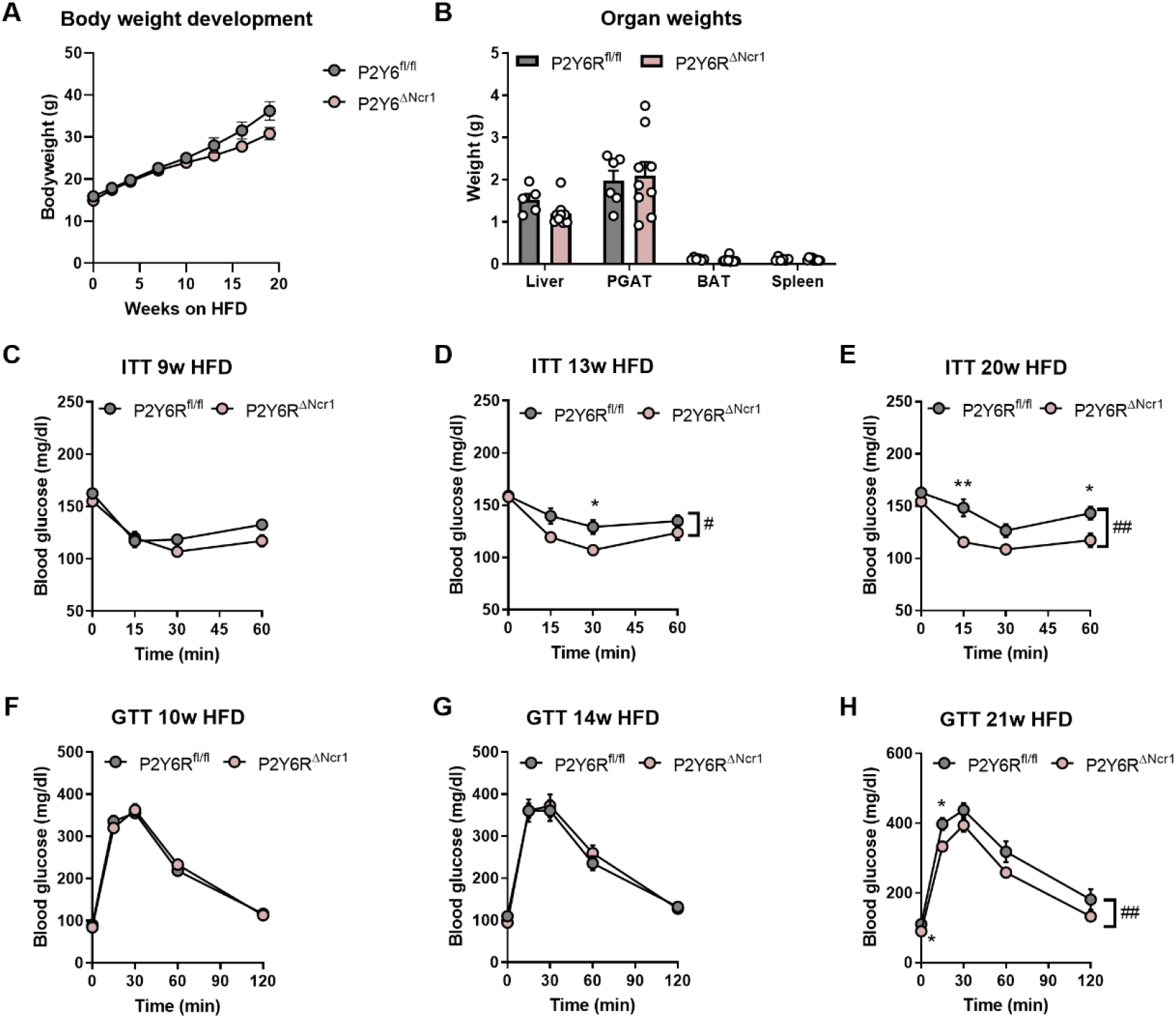
Deletion of P2Y6R from NK cells improves insulin sensitivity in female mice on HFD. **(A)** Body weight development in HFD-fed female mice (P2Y6^fl/fl^, n=16; P2Y6^ΔNcr1^, n=15) **(B)** Organ weights after 16 weeks of HFD-feeding (P2Y6^fl/fl^, n=5; P2Y6^ΔNcr1^, n=9) **(C-E)** Female transgenic mice lacking P2Y6R expression in NK cells (P2Y6^ΔNcr1^, n=15) and littermate control mice (P2Y6^fl/fl^, n=16) fed a HFD for 9 weeks **(C)**, 13 weeks **(D)** and 20 weeks **(E)** were subjected to an insulin tolerance test (ITT). **(F-H)** Glucose tolerance tests (GTT) were performed in female P2Y6^fl/fl^ (n=16) and P2Y6^ΔNcr1^ (n=15) after 10 weeks **(F)**, 14 weeks **(G)** and 21 weeks **(H)** on HFD. Statistics: (A, C-H) Two-way RM ANOVA with Sidak’s multiple comparison test, *p≤0.05, **p≤0.01; genotype: #p≤0.05, ##p≤0.01. Error bars represent SEM. (B, I-K) Unpaired two-tailed t-test, *p≤0.05. Data are represented as mean ± SEM. ○ denotes individual mice.

**Fig. 1:**
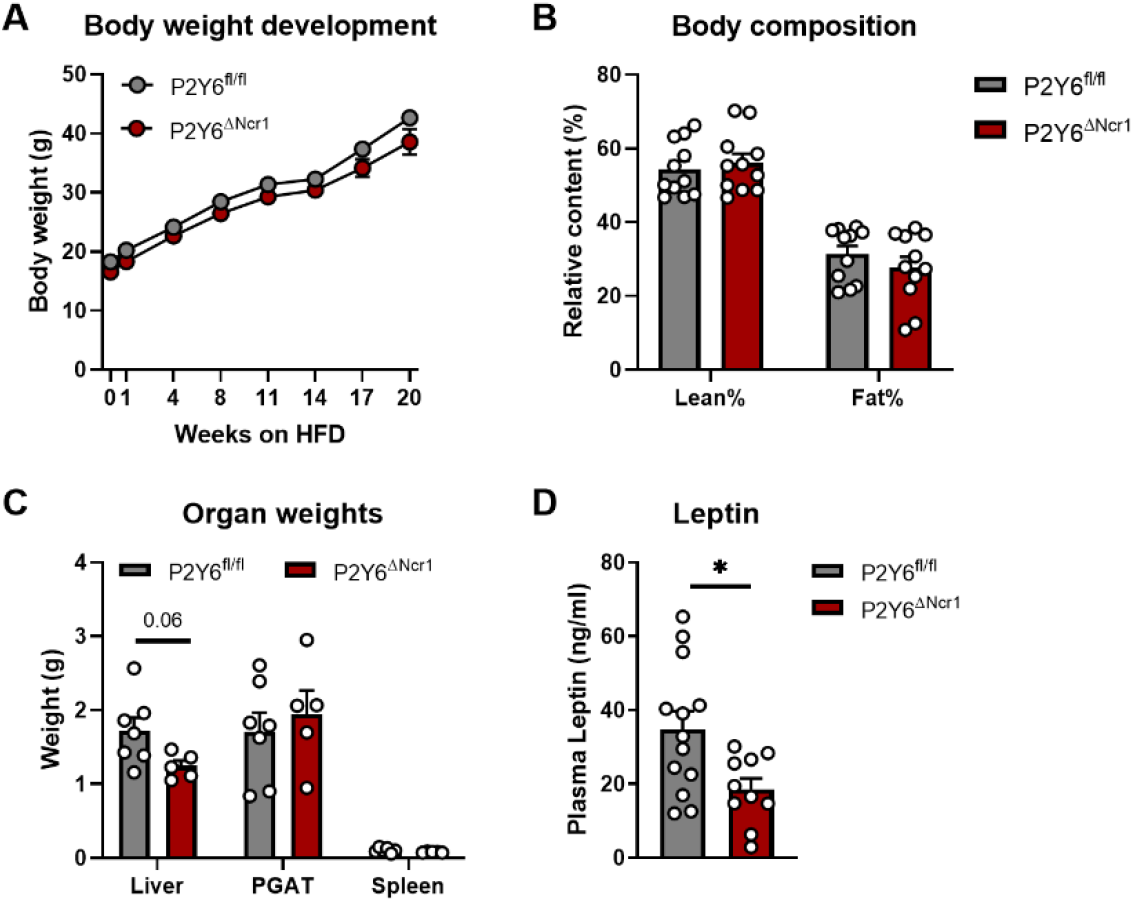
Deletion of the P2Y6R gene from NK cells does not affect energy homeostasis in mice on HFD. **(A)** Body weight development was assessed over a period of 20 weeks in transgenic mice lacking P2Y6R expression in NK cells (P2Y6^ΔNcr1^, n=11) and littermate control mice (P2Y6^fl/fl^, n=11) subjected to HFD feeding. **(B)** Body composition of P2Y6^fl/fl^ (n=11) and P2Y6^ΔNcr1^ (n=11) mice after 15-16 weeks on HFD was determined by µ-CT analysis. **(C)** Organ weights of P2Y6^fl/fl^ (n=7) and P2Y6^ΔNcr1^ (n=5) mice after 14 weeks on HFD. **(D)** Plasma leptin levels of P2Y6^fl/fl^ (n=13) and P2Y6^ΔNcr1^ (n=10) mice fed a HFD for 14 weeks and fasted for 16 hours were analyzed by ELISA. Statistics: (A) Two-way RM ANOVA with Sidak’s multiple comparison test. (B-D) Unpaired two-tailed t-test, *p≤0.05. Data are represented as mean ± SEM. ○ denotes individual mice.

### P2Y6R signaling in natural killer cells impairs insulin sensitivity in HFD mice

Next, we aimed to determine the impact of P2Y6R signaling in NK cells on glucose metabolism and insulin sensitivity. To this end, we performed insulin tolerance tests (ITT) and glucose tolerance tests (GTT) at various timepoints upon exposure to CD or HFD. Compared to controls, deletion of the P2Y6R from NK cells improved insulin sensitivity after 13 and 20 weeks of HFD feeding in both genders (Fig. 2A-C and Suppl. Fig. S3C-E) but not in male mice fed a control diet (Suppl. Fig. S5C-E). However, glucose tolerance was not affected by NK cell-specific P2Y6R deletion neither on HFD (Fig. 2D-F and Suppl. Fig. S3F-H) nor on CD (Suppl. Fig. S5F-H). While we did not observe any differences in fasted blood glucose levels between P2Y6^ΔNcr1^ mice and their control littermates on HFD (Fig. 2G), fasted insulin concentrations were reduced in P2Y6^ΔNcr1^ mice as early as after 10 weeks of HFD feeding (Fig. 2H) concomitant with a reduction of the homeostasis model assessment index for insulin resistance (HOMA-IR) (Fig. 2I). These effects were HFD-specific and could not be observed in mice on control diet (Suppl. Fig. S5I-J). Taken together, abrogation of P2Y6R-signaling from NK cells, partially protects from the development of obesity-associated insulin resistance.

**Fig. S4:**
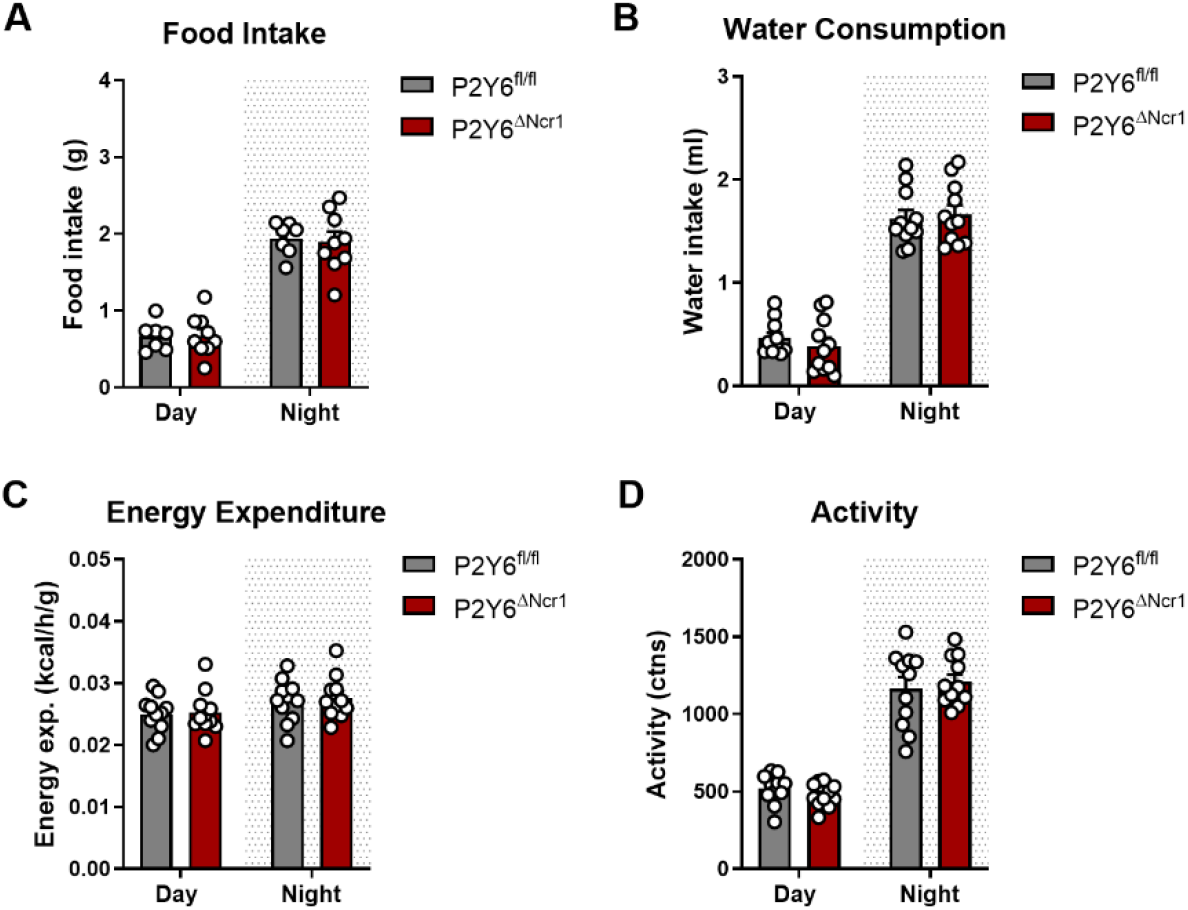
NK cell-specific deletion of P2Y6R does not affect food intake or energy expenditure in male mice on HFD. Indirect calorimetry was performed in P2Y6^fl/fl^ (n=11) and P2Y6^ΔNcr1^ (n=11) mice after 14 weeks on HFD. Data are shown as average values for day and night. **(A)** Food intake. **(B)** Water consumption. **(C)** Energy expenditure normalized to lean mass. **(D)** Average activity counts. Statistics: (A-D) Unpaired two-tailed t-test, *p≤0.05. Data are represented as mean ± SEM. ○ denotes individual mice.

**Fig. S5:**
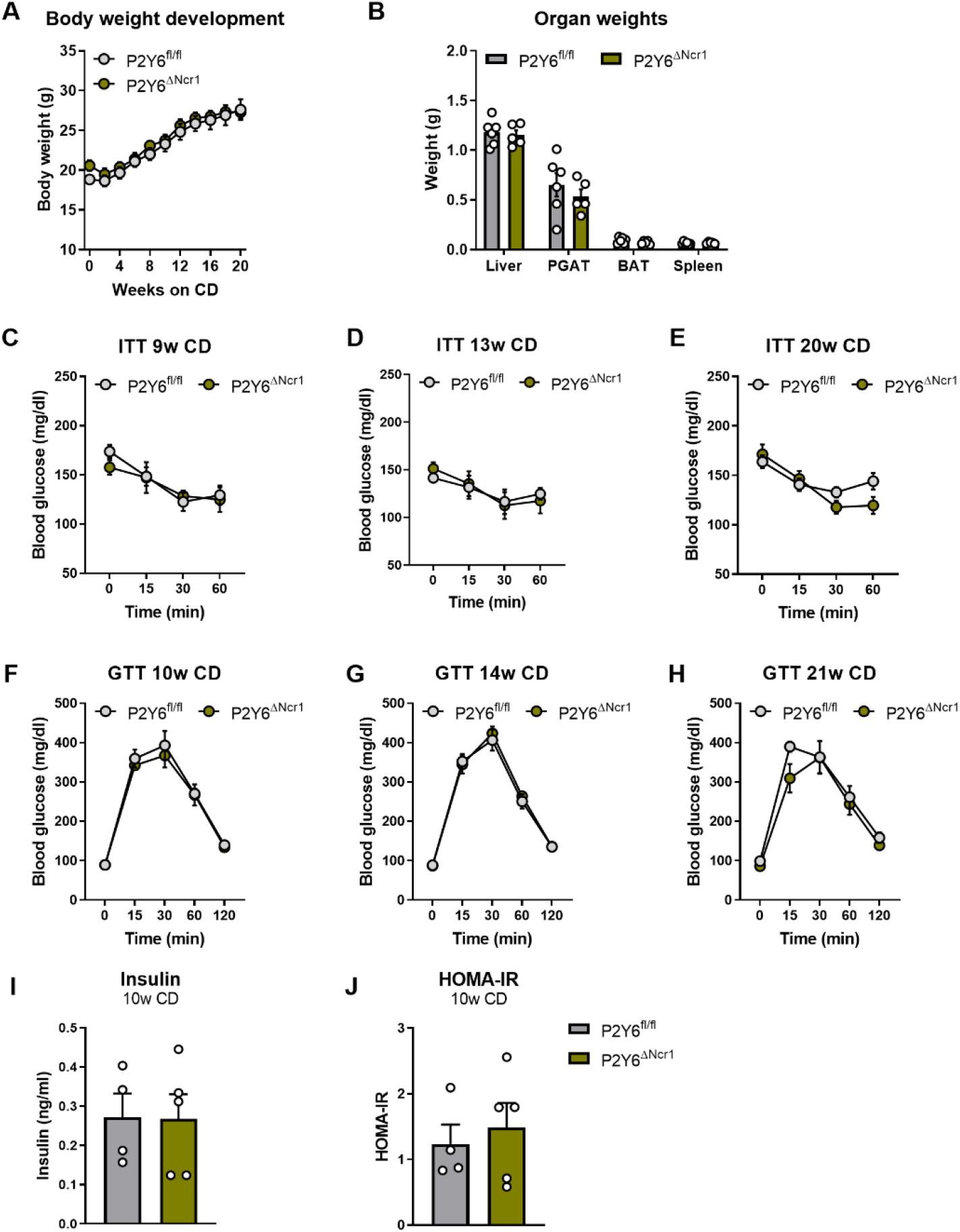
P2Y6R deletion in NK cells of control diet-fed mice does not improve insulin sensitivity. **(A)** Body weight development in control diet-fed male mice (P2Y6^fl/fl^, n=6; P2Y6^ΔNcr1^, n=5). **(B)** Organ weights after 21 weeks of CD-feeding (P2Y6^fl/fl^, n=6; P2Y6^ΔNcr1^, n=5). **(C-E)** Transgenic mice lacking P2Y6R expression in NK cells (P2Y6^ΔNcr1^, n=5) and littermate control mice (P2Y6^fl/fl^, n=6) fed a CD for 9 weeks **(C)**, 13 weeks **(D)** and 20 weeks **(E)** were subjected to an insulin tolerance test (ITT). **(F-H)** Glucose tolerance tests (GTT) were performed in P2Y6^fl/fl^ (n=6) and P2Y6^ΔNcr1^ (n=5) after 10 weeks **(F)**, 14 weeks **(G)** and 21 weeks **(H)** on CD. **(I)** Plasma insulin levels of P2Y6^fl/fl^ (n=4) and P2Y6^ΔNcr1^ (n=6) mice fed a CD for 10 weeks and fasted for 16 hours were analyzed by ELISA. **(J)** Homeostasis Model Assessment (HOMA) Index for insulin resistance (IR) was determined for P2Y6^fl/fl^ (n=4) and P2Y6^ΔNcr1^ (n=6) mice after 10 weeks on CD. Statistics: (A, C-H) Two-way RM ANOVA with Sidak’s multiple comparison test. (B, I-J) Unpaired two-tailed t-test. Data are represented as mean ± SEM. ○ denotes individual mice.

**Fig. 2:**
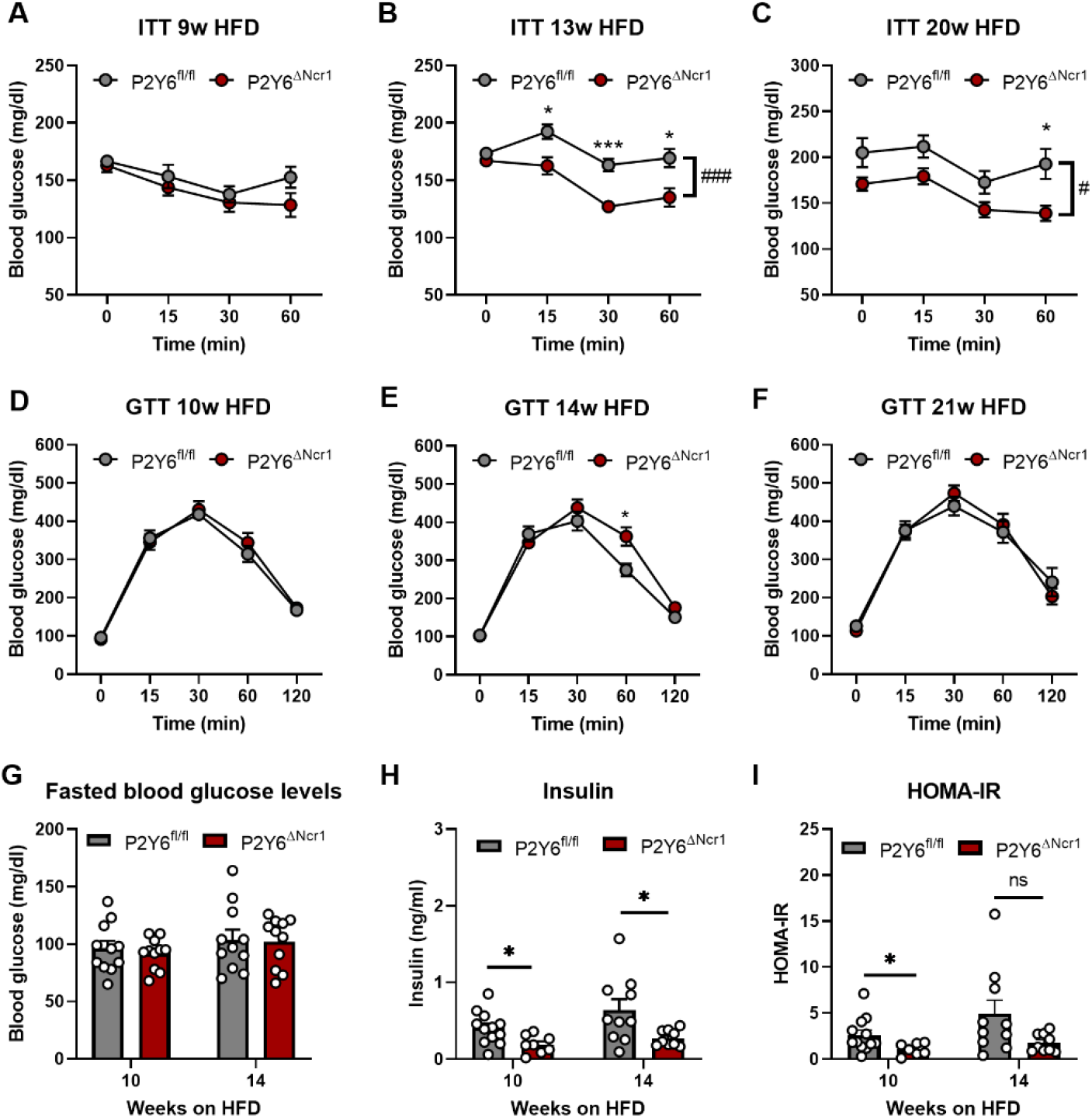
P2Y6R signaling in natural killer cells impairs insulin sensitivity in HFD-fed mice. **(A-C)** Transgenic mice lacking P2Y6R expression in NK cells (P2Y6^ΔNcr1^, n=11) and littermate control mice (P2Y6^fl/fl^, n=11) on HFD for 9 weeks **(A)**, 13 weeks **(B)** and 20 weeks **(C)** were subjected to an insulin tolerance test (ITT). **(D-F)** Glucose tolerance tests (GTT) were performed in P2Y6^fl/fl^ (n=11) and P2Y6^ΔNcr1^ (n=11) after 10 weeks **(D)**, 14 weeks **(E)** and 21 weeks **(F)** on HFD. **(G)** Blood glucose levels after 16 hour-fasting in P2Y6^fl/fl^ (n=11) and P2Y6^ΔNcr1^ (n=11) mice after 10 and 14 weeks on HFD. **(H)** Plasma insulin levels of P2Y6^fl/fl^ (n=10-12) and P2Y6^ΔNcr1^ (n=8-9) mice fed a HFD for 14 weeks and fasted for 16 hours were analyzed by ELISA. **(I)** Homeostasis Model Assessment (HOMA) Index for insulin resistance (IR) was determined for P2Y6^fl/fl^ (n=10-12) and P2Y6^ΔNcr1^ (n=7-9) mice after 10 and 14 weeks on HFD. Statistics: (A-F) Two-way RM ANOVA with Sidak’s multiple comparison test, *p≤0.05, **p≤0.01, ***p≤0.001; genotype: #p≤0.05, ###p≤0.001. Error bars represent SEM. (G-I) Unpaired two-tailed t-test, *p≤0.05. Data are represented as mean ± SEM. ○ denotes individual mice.

### NK cell-specific deletion of P2Y6R improves insulin-induced suppression of hepatic glucose production in obesity

To further dissect tissue-specific effects of P2Y6R signaling in NK cells on insulin sensitivity and glucose homeostasis, we performed hyperinsulinemic-euglycemic clamp studies in HFD-fed P2Y6^ΔNcr1^ mice and their littermate controls. Mice with NK cell-specific depletion of the P2Y6R required significantly higher glucose infusion rates (GIR) compared to their littermate controls (Fig. 3A) in order to maintain euglycemic blood glucose levels during the steady state of the clamp experiment (Fig. 3B), while exhibiting comparable levels of circulating human insulin concentrations (Fig. 3C). Quantitative assessment of insulin’s ability to suppress gluconeogenesis in both groups of mice revealed that P2Y6^ΔNcr1^ mice exhibited a clear improvement in insulińs ability to suppress hepatic glucose production (HGP) (Fig. 3D). Since hepatic glucose production is controlled both through insulin-induced changes in the expression of key enzymes of gluconeogenesis as well as through gluconeogenic substrate availability, such as fatty acids released from adipose tissue through lipolysis, we next compared the ability of insulin to regulate the concentration of plasma free fatty acid (FFA) concentrations during the clamp phase. This analysis revealed that while in obese HFD-fed control mice insulin failed to suppress circulating plasma FFA concentrations during the clamp, insulin significantly reduced circulating FFA concentrations in P2Y6^ΔNcr1^ mice (Fig. 3E).

**Fig. 3:**
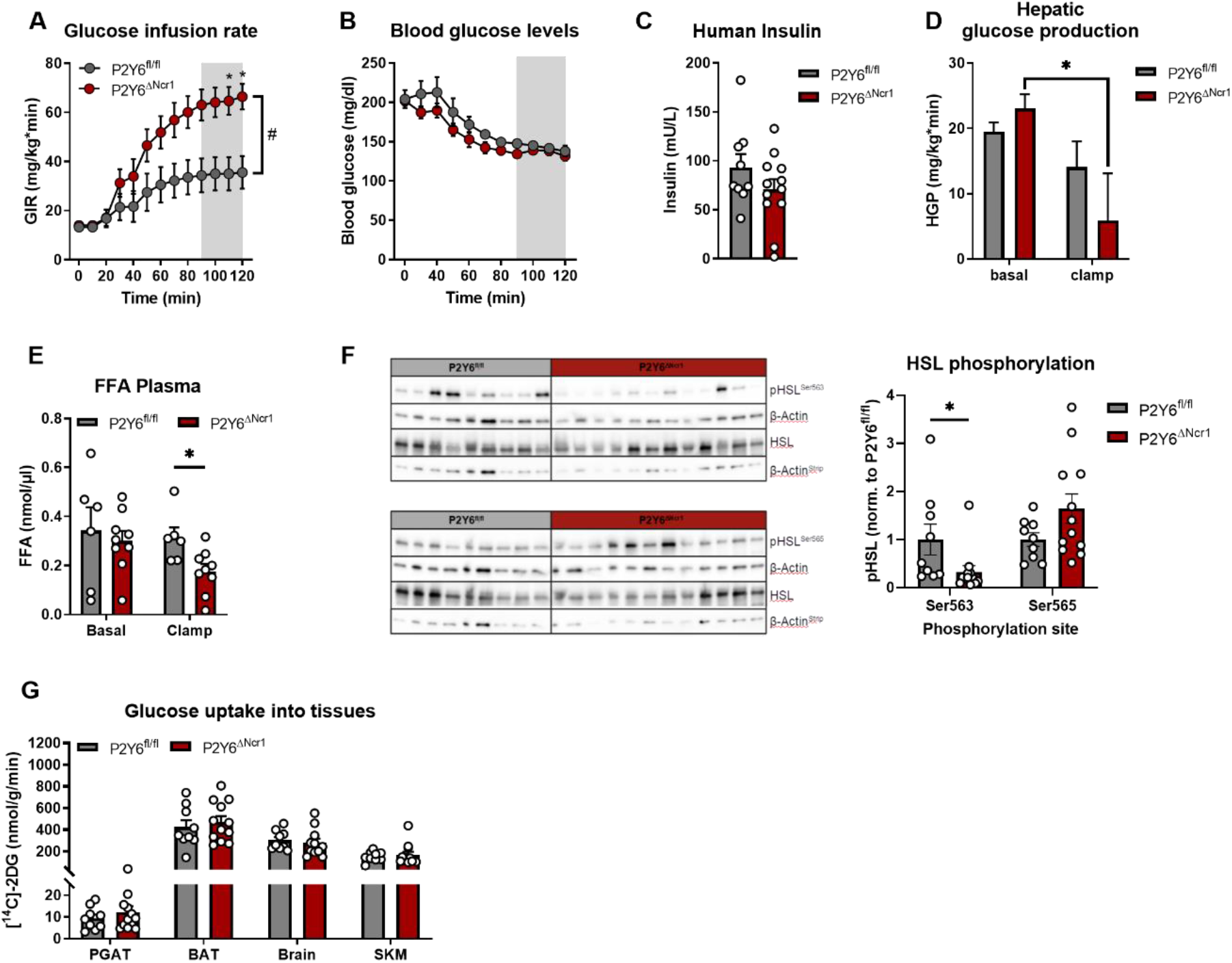
P2Y6R deletion in NK cells reduces hepatic glucose production and lipolysis in adipose tissue. Transgenic mice lacking P2Y6R expression in NK cells (P2Y6^ΔNcr1^, n=12) and littermate control mice (P2Y6^fl/fl^, n=9) were subjected to HFD feeding for 9 weeks. (A and B) Hyperinsulinemic-euglycemic clamp study **(A)** Glucose-infusion rate (GIR) required to reach euglycemic steady state glucose levels (90-120 min). **(B)** Blood glucose levels, monitored every 10 min during the experiment, reached euglycemic steady state levels at the end of the experiment. **(C)** Human insulin levels in plasma during the clamped state were determined by ELISA (P2Y6^fl/fl^, n=9; P2Y6^ΔNcr1^, n=12). **(D)** Hepatic glucose production (HGP) (P2Y6^fl/fl^, n=8; P2Y6^ΔNcr1^, n=9) and **(E)** free fatty acid (FFA) concentration in plasma (P2Y6^fl/fl^, n=6; P2Y6^ΔNcr1^, n=9) were determined at the basal and clamped state. **(F)** Organs of clamped P2Y6^fl/fl^ (n=9) and P2Y6^ΔNcr1^ (n=12) mice on HFD (9 weeks) were further investigated by western blot analysis. Representative immuno-blots for phosphorylated hormone-sensitive lipase (HSL) at Ser563 and at Ser565, total HSL and β-Actin in PGAT of clamped mice. Quantification of pHSL^Ser563^ and pHSL^Ser565^ levels was performed after normalization to β-Actin and total HSL levels. **(G)** Organ-specific glucose uptake rates during the clamped state in P2Y6^fl/fl^ (n=9) and P2Y6^ΔNcr1^ (n=12) mice. Statistics: Two-way RM ANOVA with Sidak’s multiple comparison test. *p≤ 0.05, #p (Genotype) ≤ 0.05. (B) Two-way RM ANOVA with Sidak’s multiple comparison test for steady state (90-120 min) p=0.2598. (C-H) Unpaired two-tailed t-test, *p≤.0.05. Data are represented as mean ± SEM. ○ denotes individual mice.

Lipolysis is exceptionally sensitive to the action of insulin (Jensen and Nielsen, 2007) and tightly regulated by the phosphorylation of hormone sensitive lipase (HSL) in adipose tissue (Nielsen *et al*., 2014). We found that insulin inhibits the phosphorylation of HSL at serine 563 in P2Y6^ΔNcr1^ mice (Fig. 3F), which is a main phosphorylation site known to regulate HSL-driven lipolysis (Nielsen *et al*., 2014), while phosphorylation of serine 565 only showed a mild increase in P2Y6^ΔNcr1^ mice (Fig. 3F).

However, in contrast to an enhanced ability of insulin to suppress HGP and adipose tissue lipolysis, insulin-stimulated uptake of [^14^C]-D-glucose into perigonodal adipose tissue (PGAT), brown adipose tissue (BAT), brain, and skeletal muscle (SKM) remained unaltered in P2Y6^ΔNcr1^ mice compared to their controls (Fig. 3G). Consistent with unaltered insulin-stimulated glucose uptake in these organs, also assessment of insulin-stimulated protein kinase B (AKT) phosphorylation during the clamp failed to reveal significant differences in P2Y6^ΔNcr1^ mice compared to their controls (Fig. 4). Moreover, insulin’s ability to activate AKT phosphorylation in liver did not differ between the two groups of animals (Fig. 4). Thus, deletion of P2Y6R from NK-cells in obesity appears to enhance insulin-stimulated suppression of lipolysis in PGAT and thus gluconeogenic substrate supply to liver.

**Fig. 4:**
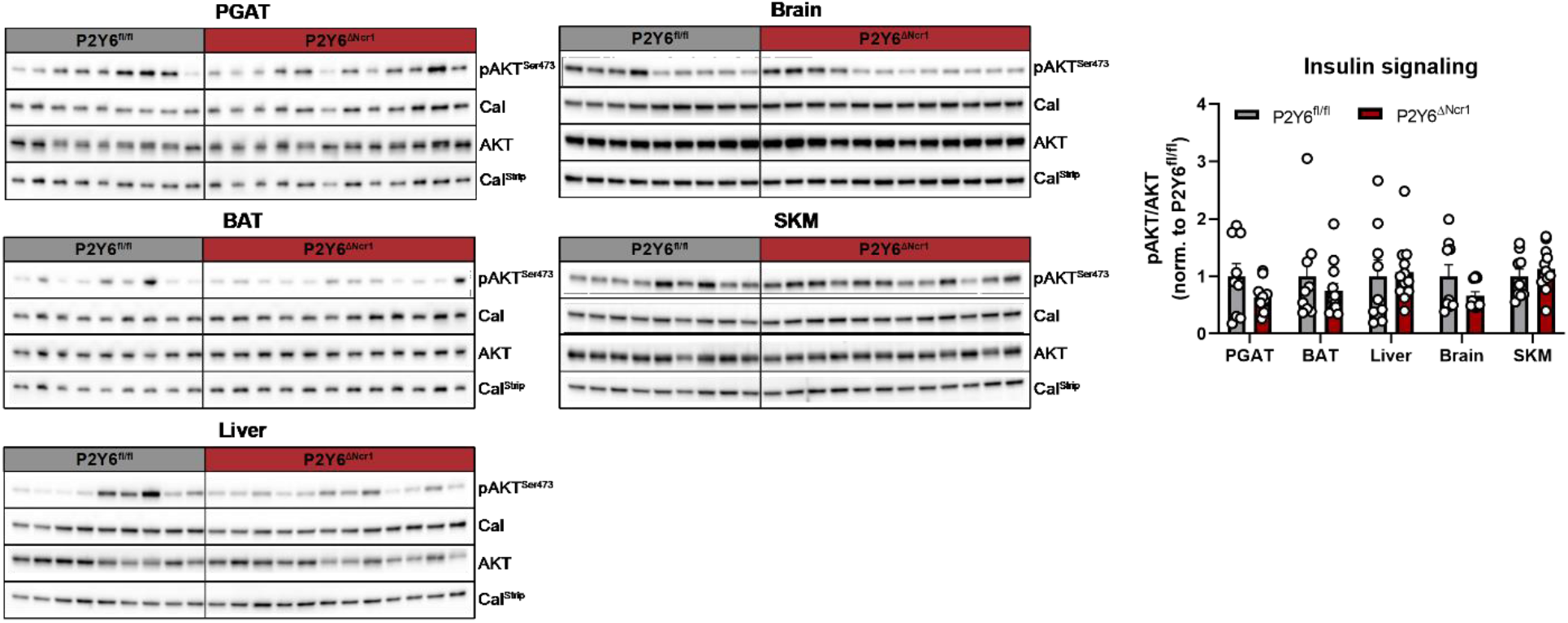
NK cell-specific deletion of P2Y6R does not alter insulin signaling in peripheral organs. Organs of clamped P2Y6^fl/fl^ (n=9) and P2Y6^ΔNcr1^ (n=12) mice on HFD (9 weeks) were further investigated by western blot analysis. Representative immuno-blots for phosphorylated AKT (Ser473), total AKT and Calnexin in PGAT, BAT, liver, brain and SKM of clamped mice. Quantification of pAKT^Ser473^ levels was performed after normalization to Calnexin and total AKT as loading controls. Statistics: Unpaired two-tailed t-test, *p≤0.05. Data are represented as mean ± SEM. ○ denotes individual mice.

### NK cell-specific deletion of P2Y6R upregulates genes associated with triglyceride homeostasis and increases mitochondrial function in liver

To further investigate the effects of NK cell-specific P2Y6R signaling on liver and adipose tissue, we compared gene expression profiles of liver and PGAT from P2Y6^ΔNcr1^ and their control littermates on HFD by mRNA deep sequencing. Differential gene expression analysis (DESeq2) identified 166 differentially expressed genes in liver and 58 genes in PGAT of P2Y6^ΔNcr1^ mice compared to their controls (Fig. 5A). To gain more functional insights based on the gene expression patterns, we subjected significantly regulated genes (cutoff p≤0.05) to gene set enrichment analyses. The three most significantly regulated gene ontologies in liver identified pathways of acylglycerol homeostasis, triglyceride homeostasis and oxidative phosphorylation (Fig. 5B), gene set enrichment analysis for regulated genes in PGAT did not yield any metabolism-related gene ontology terms (data not shown). Here, expression of lipoprotein lipase (*Lpl*), a marker gene of the GO terms for acylglycerol and triglyceride homeostasis, was shown to be specifically up-regulated in liver of P2Y6^ΔNcr1^ mice compared to controls, but not in PGAT (Fig 5C). In contrast, *Plin2* expression was significantly down-regulated in PGAT and showed a mild decrease in expression in liver (Fig. 5D).

**Fig. 5:**
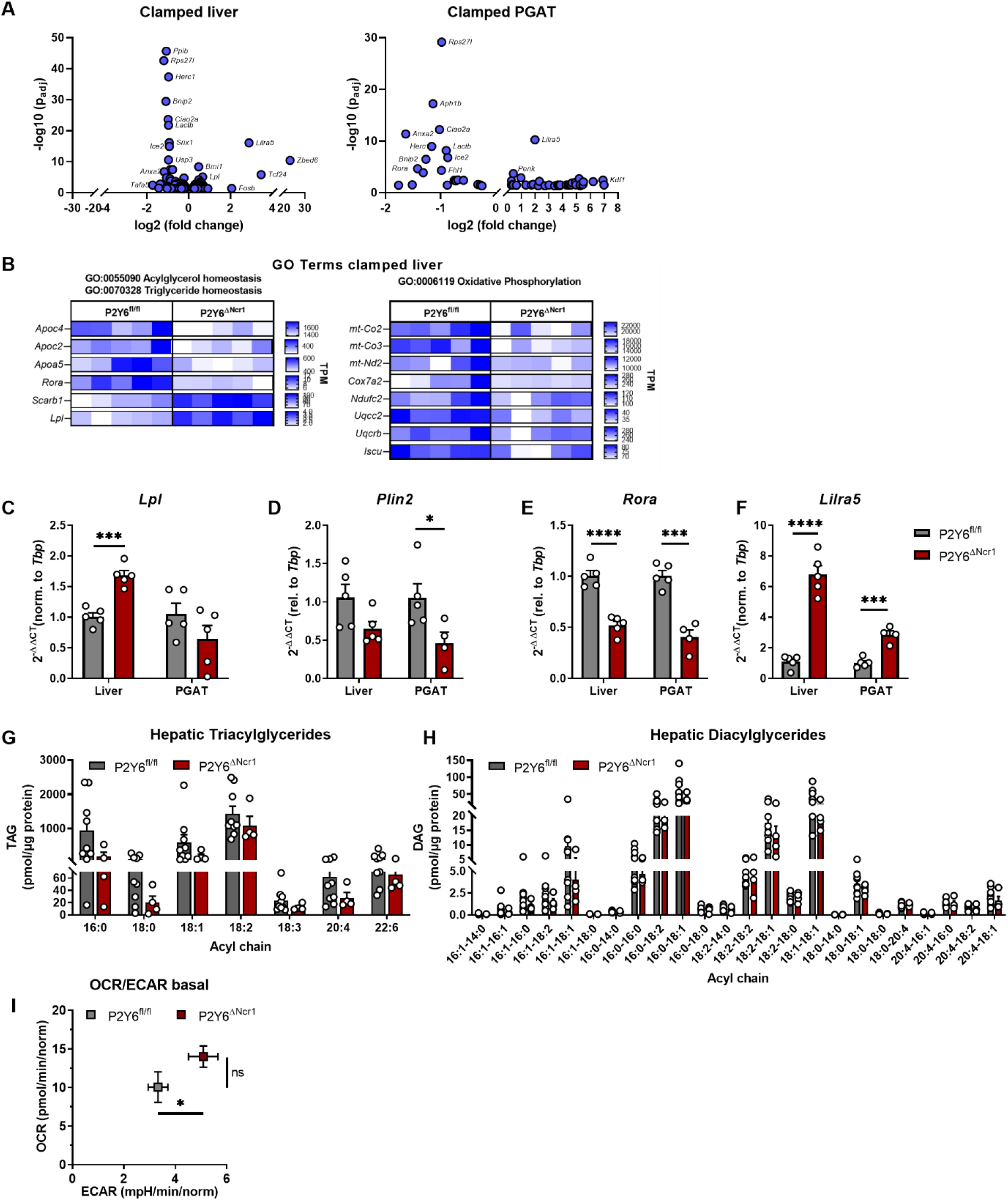
NK cell-specific deletion of P2Y6R upregulates genes associated with triglyceride homeostasis and increases mitochondrial function in liver. **(A)** Differentially expressed genes in liver (left) and PGAT (right) of clamped P2Y6^fl/fl^ (n=5) versus P2Y6^ΔNcr1^ (n=5) mice on HFD (9 weeks). Gene set enrichment analysis of transcriptomes in livers of clamped P2Y6^fl/fl^ (n=5) versus P2Y6^ΔNcr1^ (n=5) mice. Shown are the most significantly enriched gene ontology terms (GO terms) with the transcripts per million (TPM) for the GO term defining marker genes. **(C-F)** Validation of the RNASeq data by real-time PCR analysis. Expression of lipoprotein lipase (*Lpl*), RAR-related orphan receptor alpha (*Rora*), Perilipin 2 (*Plin2*) and Leukocyte immunoglobulin-like receptor subfamily A member 5 (*Lilra5*) was quantified in clamped liver and PGAT of P2Y6^fl/fl^ (n=5) and P2Y6^ΔNcr1^ (n=5) mice. Quantification was determined by the 2^-ΔΔCt^ method using the housekeeping gene TATA-binding protein (*Tbp*) for normalization. **(G-H)** Lipidomic analysis of livers from P2Y6^fl/fl^ (n=8) and P2Y6^ΔNcr1^ (n=4) mice on HFD (9 weeks). **(I)** Seahorse analysis in liver punches of P2Y6^fl/fl^ (n=5) and P2Y6^ΔNcr1^ (n=4) mice on HFD (9 weeks). Baseline oxygen consumption rate (OCR) as measure for respiration was plotted against the extracellular acidification rate (ECAR) as measure for glycolysis. Statistics: (C-I) Unpaired two-tailed t-test, *p≤0.05, **p≤0.01, ***p≤0.001, ****p≤0.0001. Data are represented as mean ± SEM. ○ denotes individual mice.

Interestingly, analysis of the overlap of commonly regulated genes in liver and PGAT revealed that 10 genes (*Lilra5*, *Rora*, *Herc1*, *Bnip2*, *Lactb*, *Ciao2a*, *Rps27l*, *Ice2*, *Anxa2*, *Vps13c*) were similarly differentially expressed in both organs (Suppl. Fig. S6). Importantly, some of them are known to be specifically expressed in immune cells and NK cells, thus raising the possibility that some of the commonly observed differences in both tissues might result from their coordinated expression changes in tissue-infiltrating NK cells. Here, only *Lilra5* expression was found to be up-regulated, whereas the expression of *Rora* and *Herc1* was down-regulated (Suppl. Fig. S6). These findings were further validated by real-time PCR analysis. Consistent with the results obtained in the mRNA sequencing experiment, *Rora* expression was significantly down-regulated in liver and PGAT of P2Y6^ΔNcr1^ mice compared to the expression in littermate controls (Fig. 5E) and *Lilra5* expression was confirmed to be up-regulated in both organs of P2Y6R knock out mice (Fig. 5F).

**Fig. S6:**
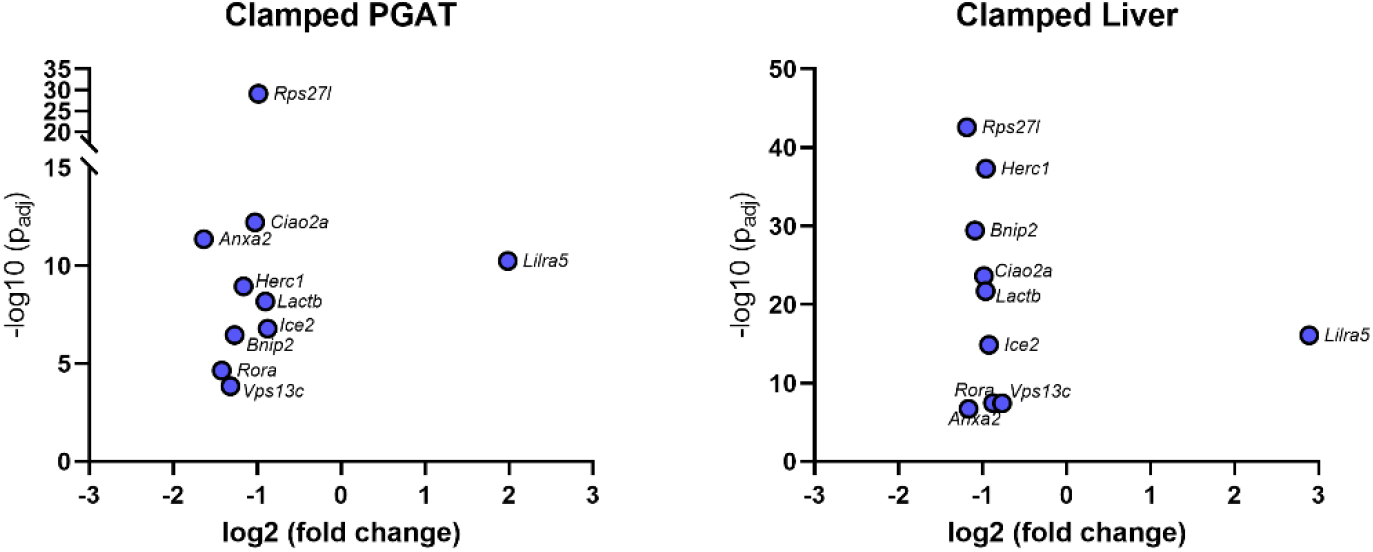
Overlap analysis of differentially expressed genes in clamped PGAT and liver reveal a NK cell-specific gene set. Differentially expressed genes (p≤0.05) in clamped PGAT and liver were compared and overlapping genes plotted as volcano plots.

As triglyceride (TG) homeostasis was affected by NK cell-specific deletion of P2Y6R, we conducted lipidomic analyses of liver samples from mice fed a HFD for 9 weeks. These analyses revealed no significant overall changes in hepatic TG (Fig. 5G) and diacylglyceride (DAG) contents (Fig. 5H), however some specific lipid species showed a trend towards reduction in liver of P2Y6^ΔNcr1^ compared to controls (C16:0, C18:0 TGs, C16:0-C18:1, C18:1-C18:1 DAGs).

Since hepatic lipid homeostasis is closely linked to mitochondrial function in liver, we next investigated the rates of oxidative capacity and extracellular acidification rates (ECAR) in micro-punches from liver of HFD-fed P2Y6^ΔNcr1^ mice and their control littermates. This analysis revealed a significant upregulation of the ECAR (Fig. 5I), indicating an increase in glycolysis, while the oxygen consumption rate (OCR) was only slightly but not significantly improved in liver of HFD-fed P2Y6^ΔNcr1^ mice compared to their control littermates (Fig. 5I).

Collectively, NK cell-specific P2Y6R signaling appears to affect triglyceride metabolism and mitochondrial function in liver, while coordinately regulated differences in gene expression between liver and adipose tissue possibly arise from NK cell intrinsic changes in gene expression in tissue-infiltrating NK cells.

### NK cell specific deletion of P2Y6R alters tissue-infiltration of immune cells and expression of chemokines

NK cells represent a heterogeneous innate lymphocyte cell population with organ- and microenvironment-specific expression of cytotoxic molecules, receptors and transcription factors (Crinier *et al*., 2018; Zhao *et al*., 2020). As the overlap analysis of the transcriptional changes from adipose tissue and liver revealed a common set of regulated, possibly immune cell-specific genes, we next assessed the transcriptome of isolated NK cells. To this end, NK cells were isolated from adipose tissue and liver of HFD-fed P2Y6^ΔNcr1^ and P2Y6^fl/fl^ mice by flow cytometry cell sorting based on the expression of the NK cell-specific surface markers NK1.1 and Ncr1 (Suppl. Fig. S7). DESeq2 analysis of the transcriptome of bulk-sequenced NK cells resulted in 59 differentially expressed genes in adipose tissue-derived NK cells (Fig. 6A) and 78 genes in hepatic NK cells (Fig. 6B). Gene set enrichment analyses of the differentially expressed genes (cutoff p≤0.05) in adipose tissue-derived NK cells identified pathways of lymphocyte chemotaxis, lymphocyte migration and neutrophil migration (Fig. 6C), whereas pathways of protein folding, and chaperone cofactor-dependent protein folding were detected in liver-derived NK cells (Fig. 6D).

**Fig. S7:**
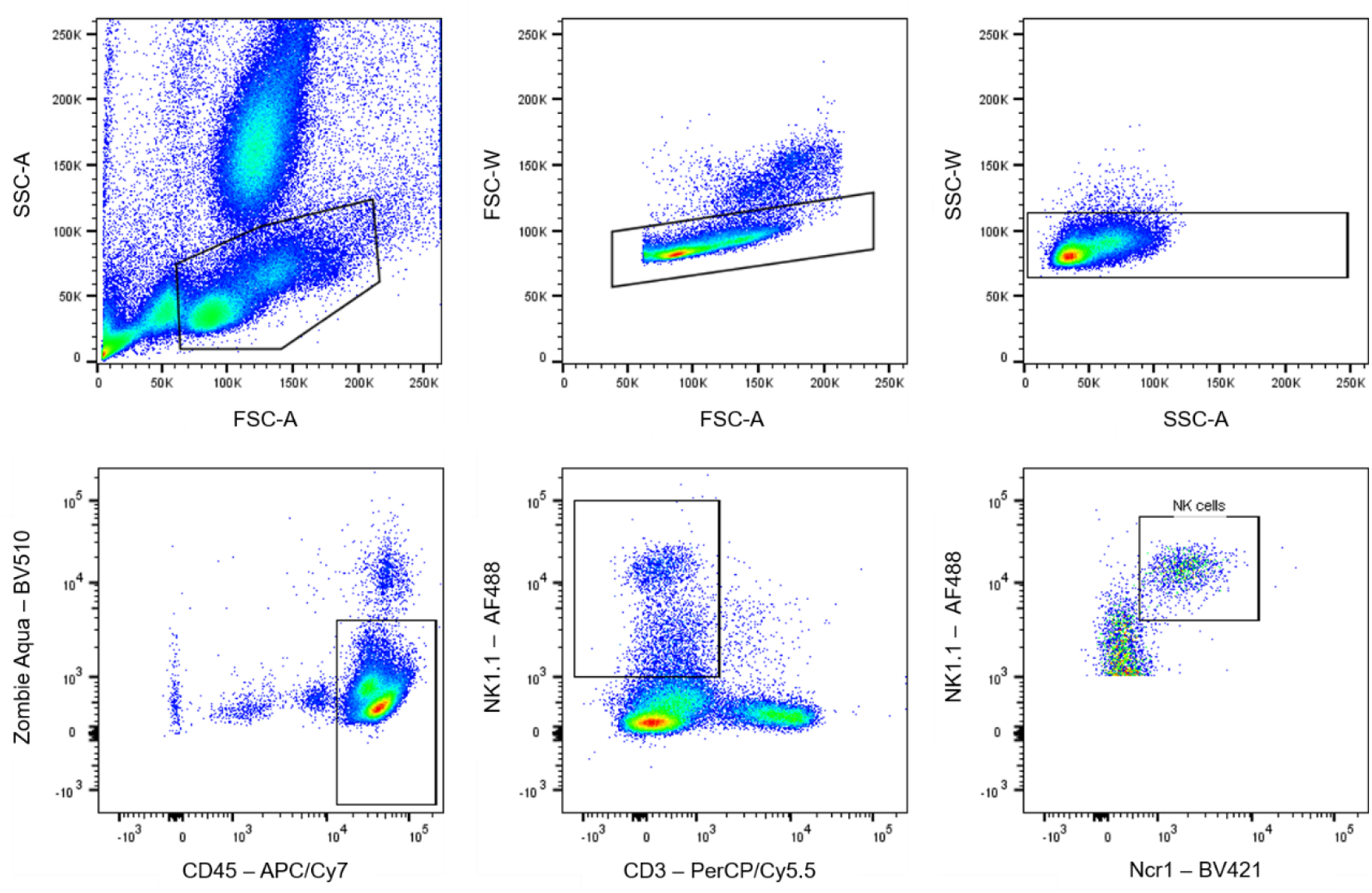
NK cells were sorted from liver, PGAT and spleen of male mice on HFD by flow cytometry. NK cells were gated as single/viable/CD45^+^/CD3^-^/NK1.1^+^/Ncr1^+^ cells.

**Fig. 6:**
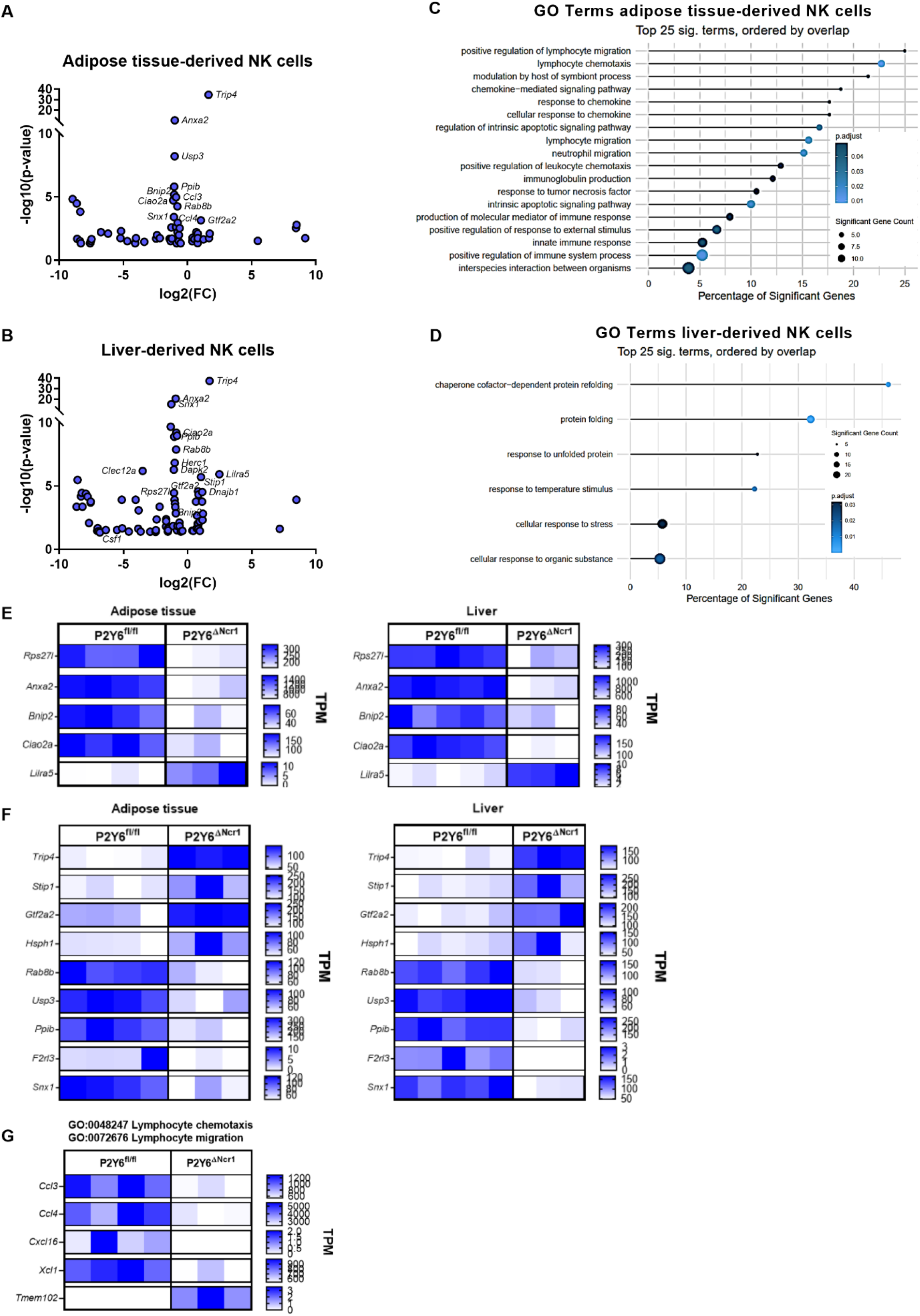
Adipose tissue-derived NK cells from P2Y6^ΔNcr1^ mice show reduced expression of chemokines. **(A-B)** Transcriptomic analysis of NK cells (defined as single/viable/CD45^+^/CD3^-^ /NK1.1^+^/Ncr1^+^ cells and sorted by flow cytometry) from adipose tissue **(A)** and liver **(B)** of P2Y6^fl/fl^ (n=4-5) and P2Y6^ΔNcr1^ (n=3) mice on HFD (15 weeks). Volcano-plots of differentially expressed genes with p≤0.05. **(C-D)** GO term analysis of differentially expressed genes with p≤0.05 in adipose tissue-derived NK cells **(C)** and **(D)** hepatic NK cells. **(E)** TPM plots of commonly differentially expressed genes in NK cells isolated from PGAT and from liver. These genes have also been found to be differentially expressed in liver and PGAT of clamped mice. **(F)** TPM plots of genes differentially expressed in NK cells from both origins, PGAT (left) and liver (right). **(G)** TPM plots of GO term defining genes differentially expressed in adipose tissue-derived NK cells of P2Y6^fl/fl^ (n=4) and P2Y6^ΔNcr1^ (n=3) mice.

Consistent with our previous findings in liver and adipose tissue, we also found genes of the presumably NK cell-specific gene set to be differentially expressed in isolated NK cells of both tissue origins (Fig. 6E). This supports our hypothesis that similar differences in gene expression between liver and adipose tissue possibly arise from NK cell intrinsic changes in gene expression of tissue-infiltrating NK cells. Besides this first NK cell-specific gene set, we found a second set of overlapping genes in isolated NK cells derived from adipose tissue and liver. This gene set includes the up-regulated genes *Trip4*, *Stip1*, *Gtf2a2* and *Hsph1* and the down-regulated genes *Rab8b*, *Usp3*, *Ppib, F2rl3* and *Snx1* (Fig. 6F)*. Usp3*, *Ppib* and *Snx1* have also been found to be differentially expressed in liver and *Rab8b* in adipose tissue of clamped P2Y6^ΔNcr1^ mice when compared to their littermate controls (data not shown).

Notably, in adipose tissue-derived NK cells the gene expression of the chemokines *Ccl3*, *Ccl4*, *Xcl1* and *Cxcl16* (Fig. 6G) was found to be down-regulated in parallel with the down-regulation of *Csf1* expression in hepatic NK cells (Fig. 6B). As these chemokines are involved in the recruitment and activation of immune cells, we aimed to elucidate the impact of this P2Y6R-dependent and NK cell-specific down-regulation on the infiltration of immune cells into different organs. In line with this reduced expression of chemokines in NK cells, we observed a significant reduction of NK cell numbers in liver and spleen, a decrease in NKT cell numbers in PGAT and a reduction of macrophages in spleen of P2Y6^ΔNcr1^ mice compared to control (Fig. 7A). In addition, infiltration of MAC2 positive macrophages into adipose tissue was reduced in mice with NK cell-specific P2Y6R deletion after prolonged HFD-feeding (Suppl. Fig. S8A), concomitant with a mild but non-significant reduction in the expression of inflammatory cytokines and chemokines (Suppl. Fig. S8B). On the contrary, expression of inflammatory markers in liver of mice fed a HFD for 22 weeks were not significantly altered between genotypes (Suppl. Fig. S8C).

**Fig. S8:**
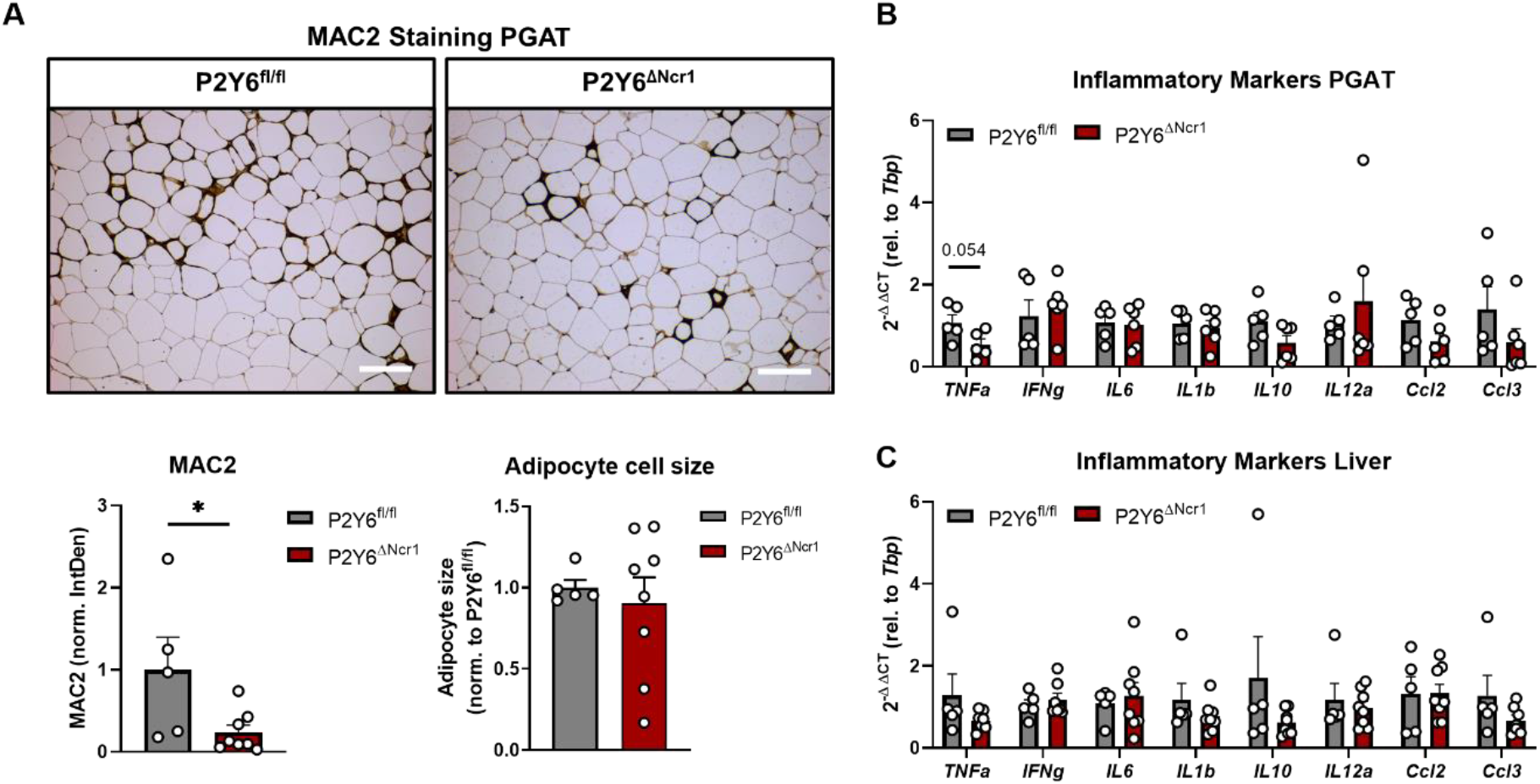
NK cell-specific deletion of P2Y6R reduces macrophage infiltration into adipose tissue in mice on HFD. Transgenic mice (P2Y6^ΔNcr1^, n=8) and their control littermates (P2Y6^fl/fl^, n=5) were fed a HFD for 22 weeks. **(A)** Immunohistochemical analysis of MAC2 positive macrophages in PGAT of P2Y6^ΔNcr1^ (n=8) and P2Y6^fl/f^ mice (n=5). Scale bars 100 µm. **(B-C)** Real-time PCR analysis of PGAT **(B)** and liver **(C)** from P2Y6^ΔNcr1^ (n=6-8) and their control littermates (P2Y6^fl/fl^, n=5). Statistics: Unpaired two-tailed t-test, *p≤0.05. Data are represented as mean ± SEM. ○ denotes individual mice.

**Fig. 7:**
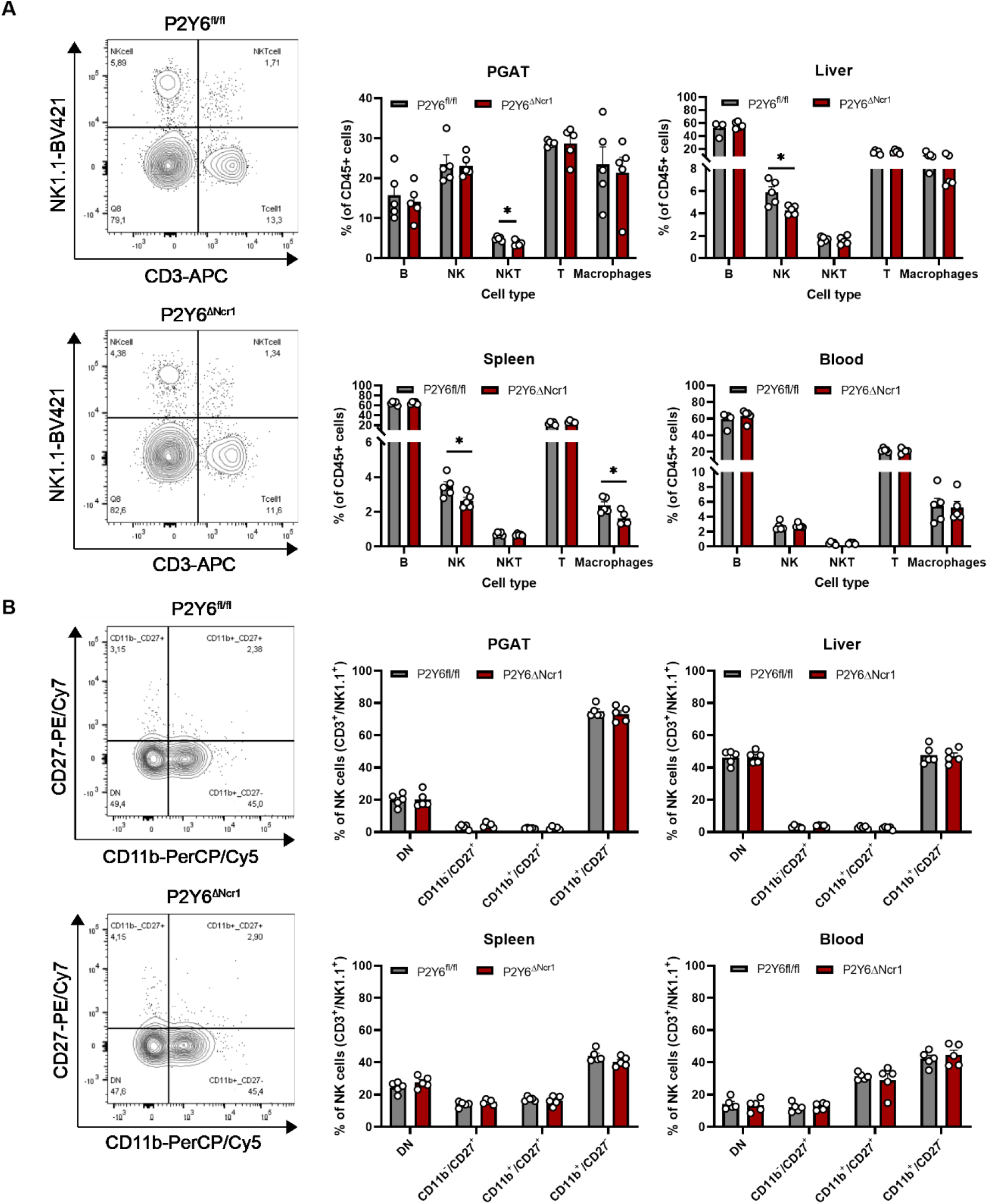
P2Y6^ΔNcr1^ mice exhibit reduced immune cell infiltration into peripheral organs. **(A)** Representative flow cytometry analysis plots and quantification of immune cell subsets in PGAT, liver, spleen and blood of P2Y6^fl/fl^ (n=5) and P2Y6^ΔNcr1^ (n=5) mice fed a HFD for 14 weeks. B cells were gated as single/viable/CD45^+^/CD3^-^/CD19^+^ cells, NK cells as single/viable/CD45^+^/CD3^-^/NK1.1^+^ cells, NKT cells as single/viable/CD45^+^/CD3^+^/NK1.1^+^ cells, T cells as single/viable/CD45^+^/CD3^+^/NK1.1^-^ cells and macrophages as single/viable/CD45^+^/CD11b^+^/F4/80^+^ cells. **(B)** Representative FACS plots and quantification of the maturation status of NK cells in PGAT, liver, spleen and blood of P2Y6^fl/fl^ (n=5) and P2Y6^ΔNcr1^ (n=5) mice fed a HFD for 14 weeks. NK cells were gated as single/viable/CD45^+^/CD3^-^/NK1.1^+^ cells and further subdivided into double negative (DN) (CD11b^-^/CD27^-^), immature (CD11b^-^/CD27^+^), intermediate (CD11b^+^/CD27^+^) and mature (CD11b^+^/CD27^-^) NK cells. Statistics: (A, B) Unpaired two-tailed t-test, *p≤0.05. Data are represented as mean ± SEM. ○ denotes individual mice.

Depending on their maturation state, NK cells express a distinct repertoire of cytotoxins and cytokines (Crinier *et al*., 2018). As we observed a reduction in chemokine expression in P2Y6^ΔNcr1^-derived NK cells, we investigated the impact of P2Y6R signaling on the maturation state of NK cells based on the surface expression of CD27 and CD11b. Quantification of the different maturation subsets of NK cells by flow cytometry analysis revealed that the majority of NK cells in adipose tissue and liver belong to the mature subset (CD11b^+^/CD27^-^), whereas in spleen and blood also a notable proportion of immature (CD11b^-^/CD27^+^) and intermediate (CD11b^+^/CD27^+^) NK cells could be found (Fig. 7B). However, abrogation of P2Y6R signaling did not affect the overall maturation state in NK cells (Fig. 7B).

In summary, P2Y6R signaling in NK-cells of HFD-fed mice likely increases the expression of chemokines favoring the infiltration of immune cells into metabolically relevant organs, while it at the same time drives NK cell intrinsic changes in the expression of specific gene sets that might serve as possible markers for tissue-infiltrating NK cells.

### Abrogation of P2Y6R-signaling in macrophages does not protect from HFD-induced insulin resistance

Since P2Y6Rs are also expressed in macrophages (Bar *et al*., 2008; Lattin *et al*., 2008) where its’ activation increases the expression of inflammatory (Bar *et al*., 2008) or regulatory cytokines (Oishi *et al*., 2016), we also investigated the role of P2Y6R signaling in macrophages in the context of obesity and the development of metaflammation-induced insulin resistance. Therefore, we crossed mice carrying a loxP-flanked P2Y6R gene (Jain *et al*., 2020) with LysM-Cre mice (Clausen *et al*., 1999). Gene expression analysis of cultured bone marrow-derived macrophages (BMDM) isolated from control (P2Y6^fl/fl^) or macrophage-specific P2Y6R-deleted (P2Y6^ΔLysM^) mice confirmed successful and specific reduction of P2Y6R expression in macrophages (Suppl. Fig. S9A). As obesity development is accompanied by a macrophage switch from M2-to M1-polarized macrophages in adipose tissue (Castoldi *et al*., 2016), we aimed to elucidate the role of macrophage polarization on P2Y6R expression. Here, we found a more than three-fold upregulation of P2Y6R expression in M2-polarized compared to M1-polarized bone marrow-derived macrophages (Suppl. Fig. S9B).

**Fig. S9:**
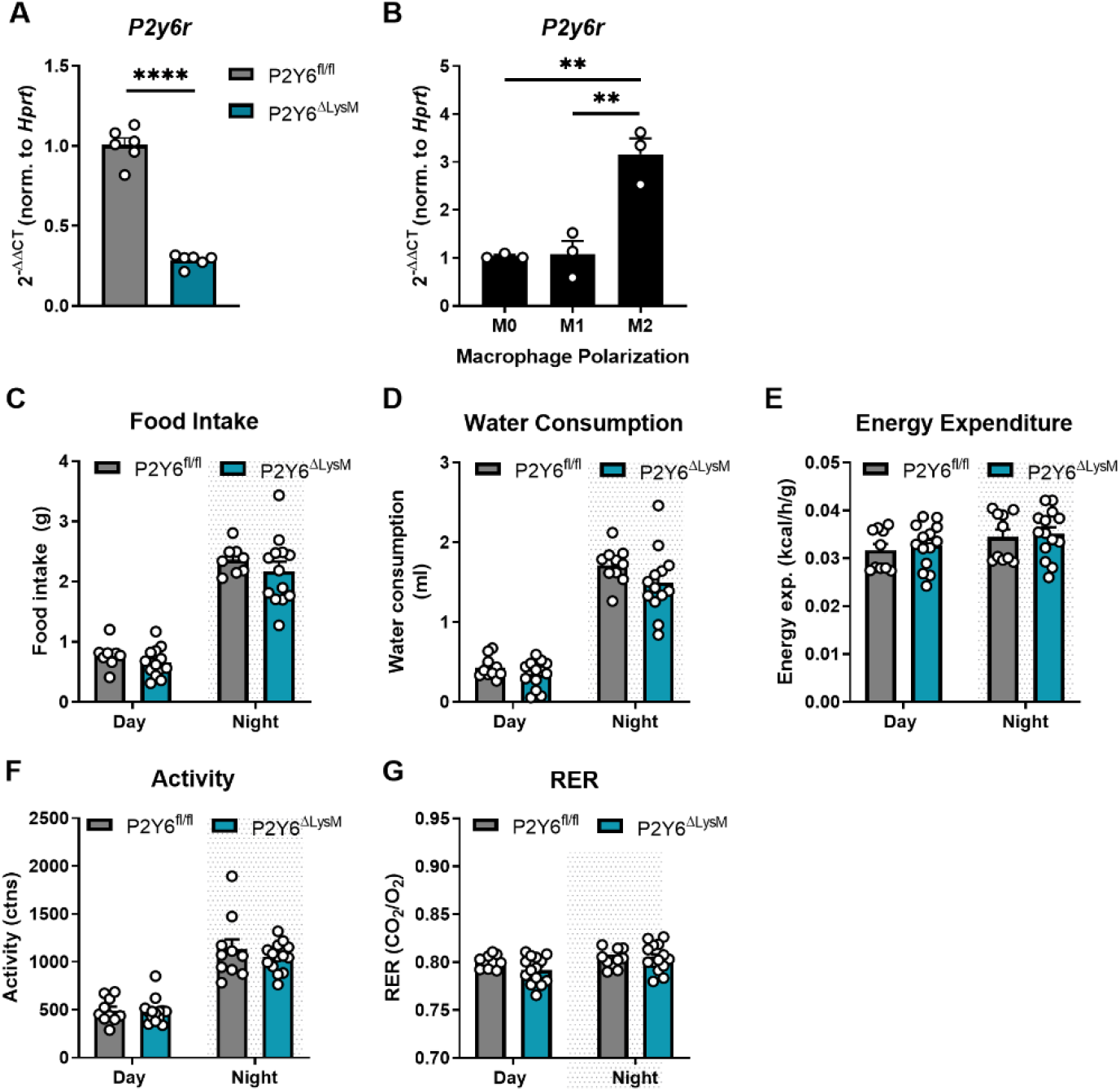
P2Y6R deletion in macrophages does not alter food intake or energy expenditure. **(A)** Bone-marrow-derived macrophages were isolated from P2Y6^fl/fl^ and P2Y6^ΔLysM^ mice and cultured for 7 days. P2Y6R expression was verified to be reduced in BMDMs of P2Y6^ΔLysM^ mice (n=6) compared to control (P2Y6^fl/fl^, n=6). Data from two independent experiments. ○ denotes individual mice. **(B)** BMDM derived from C57Bl6 mice were cultured for 7 days and polarized into M1 or M2 macrophages. P2Y6R expression in M1 and M2 BMDMs was determined by real-time PCR and normalized to the expression in unpolarized (M0) BMDMs. Data pooled from three independent experiments. ○ denotes an individual experiment. **(C-G)** Indirect calorimetry of mice with macrophage-specific deletion of P2Y6R (P2Y6^ΔLysM^, n=14) and their control littermates (P2Y6^fl/fl^, n=10) after 18 weeks on HFD. ○ denotes individual mice. Statistics: (A) Unpaired two-tailed t-test. ****p≤0.0001 (B) Two-way RM ANOVA with Sidak’s multiple comparison test. **p≤0.01; (C-G) Unpaired two-tailed student’s t-test. Data are represented as mean ± SEM.

We then investigated the impact of macrophage-specific P2Y6R signaling on the metabolic phenotype of HFD-fed mice. Mice with macrophage-specific deletion of P2Y6R showed the same body weight development as their control littermates (Fig. 8A) as well as no differences in body fat content (Fig. 8B). In parallel, investigations of insulin sensitivity and glucose tolerance after 10, 14 or 20 weeks of HFD-feeding did not reveal any significant differences between P2Y6^ΔLysM^ and P2Y6^fl/fl^ mice (Fig. 8C-H). This was in line with results from indirect calorimetry in mice fed a HFD for 18 weeks. Analyses of food intake, water consumption, energy expenditure, total activity or respiratory exchange rate (RER) did not reveal any differences between P2Y6^ΔLysM^ and P2Y6^fl/fl^ mice (Suppl. Fig. S9C-G). Also, insulin and leptin levels in plasma of 16 hours-fasted mice did not differ between genotypes concomitant with unaltered HOMA-IR (Fig. 8I-K).

**Fig. 8:**
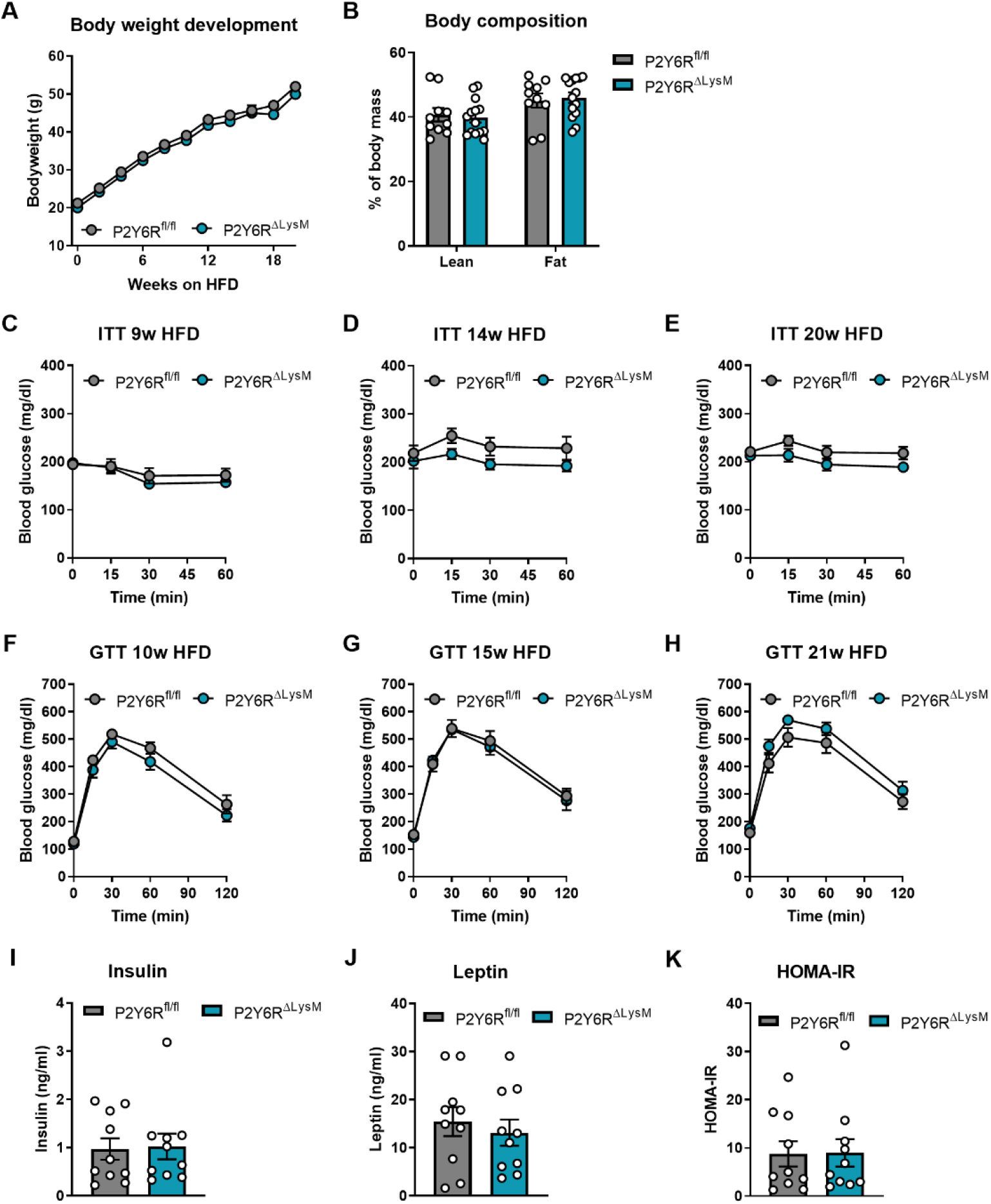
Deletion of P2Y6R in macrophages does not affect glucose metabolism in mice on HFD. **(A)** Body weight development was assessed over a period of 20 weeks in transgenic mice lacking P2Y6R expression in macrophages (P2Y6^ΔLysM^, n=14) and littermate control mice (P2Y6^fl/fl^, n=10) subjected to high-fat diet (HFD) feeding. **(B)** Body composition was investigated by µCT measurements after 19 weeks on HFD (P2Y6^fl/fl^, n=10; P2Y6^ΔLysM^, n=14) **(C-E)** Transgenic mice lacking P2Y6R expression in macrophages (P2Y6^ΔLysM^, n=14) and littermate control mice (P2Y6^fl/fl^, n=10) fed a HFD for 9 weeks **(C)**, 13 weeks **(D)** and 20 weeks **(E)** were subjected to an insulin tolerance test (ITT). **(F-H)** Glucose tolerance tests (GTT) were performed in P2Y6^fl/fl^ (n=10) and P2Y6^ΔLysM^ (n=14) after 10 weeks **(F)**, 14 weeks **(G)** and 21 weeks **(H)** on HFD. **(I)** Plasma insulin levels of P2Y6^fl/fl^ (n=10) and P2Y6^ΔLysM^ (n=10) mice fed a HFD for 10 weeks and fasted for 16 hours were analyzed by ELISA. **(J)** Blood leptin levels after 16 hour-fasting in P2Y6^fl/fl^ (n=10) and P2Y6^ΔLysM^ (n=10) mice after 10 weeks on HFD. **(K)** Homeostasis Model Assessment (HOMA) Index for insulin resistance (IR) was determined for P2Y6^fl/fl^ (n=10) and P2Y6^ΔLysM^ (n=10) mice after 10 weeks on HFD. Statistics: (A, C-H) Two-way RM ANOVA with Sidak’s multiple comparison test. (B, I-K) Unpaired two-tailed t-test, *p≤0.05. Data are represented as mean ± SEM. ○ denotes individual mice.

Since UDP levels are increased in adipocytes (Suppl. Fig. S1H) where it can serve as a “find me” signal triggering macrophages to eliminate necrotic cells, such as hypertrophic and necrotic adipocytes (Cinti *et al*., 2005), we examined the infiltration of macrophages into adipose tissue of HFD-fed mice by immunohistochemistry. However, we did not find any differences in the amount of MAC2 positive macrophages between in P2Y6^ΔLysM^ and P2Y6^fl/fl^ mice (Suppl. Fig. S10). In line with the overall inconspicuous phenotype of P2Y6^ΔLysM^ mice, we also did not observe any differences in lipid accumulation in liver between genotypes (Suppl. Fig. S10).

**Fig. S10:**
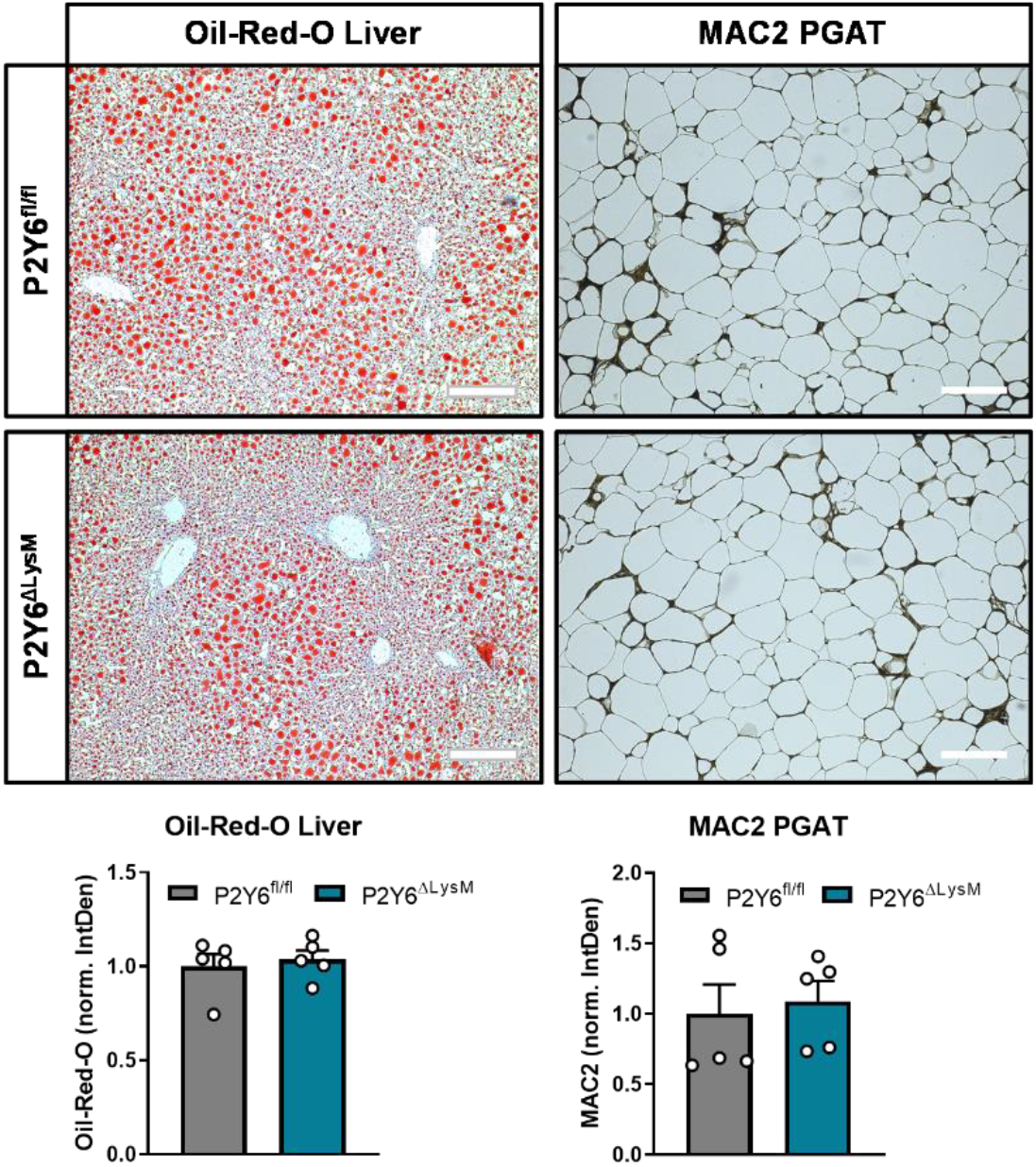
P2Y6R signaling in macrophages does not affect metaflammation in PGAT of mice on HFD. Transgenic mice (P2Y6^ΔLysM^, n=5) and their control littermates (P2Y6^fl/fl^, n=5) were fed a HFD for 22 weeks. **(left)** Oil-Red-O staining of liver tissue. Scale bars 100 µm. **(right)** Immunohistochemical analysis of MAC2 positive macrophages in PGAT. Scale bars 100 µm. Statistics: Unpaired two-tailed t-test. Data are represented as mean ± SEM. ○ denotes individual mice.

In conclusion, we demonstrated here that P2Y6R-signaling in macrophages does not play a major role in the context of obesity-induced insulin resistance and metaflammation.

## Discussion

Obesity and its’ co-morbidities are often associated with a chronic low-grade inflammation, termed “metaflammation”, characterized by increased levels of inflammatory cytokines in serum and infiltration of immune cells into adipose tissue. One immune cell type gaining increasing attention in the context of obesity and glucose metabolism are natural killer cells, a type of innate lymphoid cell (ILC) responsible for surveillance and extinction of virus-infected and tumor cells. In a previous study we reported that P2Y6R expression is up-regulated in NK cells derived from PGAT of HFD-fed mice (Theurich *et al*., 2017). P2Y6R-signaling has been reported to modulate the production of pro-inflammatory cytokines and chemotaxis of immune cells (Chen *et al*., 2014) and to be a relevant regulator of food intake and glucose metabolism in obesity (Steculorum *et al*., 2015; Steculorum *et al*., 2017).

Here, we demonstrated that UDP and uridine levels are elevated upon HFD feeding. The increase of UDP levels in plasma, adipocytes and adipose tissue-derived SVF after 12 weeks of HFD-feeding coincidences with the development of insulin resistance in wildtype mice. Mice with a NK cell-specific abrogation of P2Y6R signaling were protected from this diet-induced insulin resistance and showed a reduction in the expression of pro-inflammatory chemo- and cytokines in adipose tissue-derived NK cells. NK cells in adipose tissue have been found to induce the infiltration of macrophages mainly by an increased production and release of IFNγ (Wensveen *et al*., 2015), primarily explaining why genetic depletion of NK cells improves adipose tissue inflammation and insulin resistance (Lee *et al*., 2016). As we did not observe any alterations in the numbers of NK cells but of macrophages and NKT cells in adipose tissue upon NK cell-specific P2Y6R deletion, we rather postulated that the observed reduction in adipose tissue inflammation and associated improvement in insulin sensitivity might be due to a reduced release of inflammatory cytokines from NK cells. And indeed, we found that adipose tissue-derived NK cells showed a decreased gene expression of inflammatory molecules such as *Ccl3*, *Ccl4*, *Xcl1* and *Cxcl16*, presumably leading to an attenuation of obesity-induced metaflammation.

Obesity-induced metaflammation is a trigger for the development of insulin resistance which is in turn correlated with increased lipolysis in adipose tissue leading to a rise in FFA levels in serum of obese mice and humans (Arner and Rydén, 2015). These FFAs serve as energy source and can, after β-oxidation, be transformed to ATP by oxidative phosphorylation in mitochondria. Here, we show that NK cell-specific disruption of P2Y6R-signaling improves insulin sensitivity in mice on HFD, most prominently reflected by an insulin-induced reduction of lipolysis in adipose tissue and repression of glucose production in liver. A causal linkage between insulin suppression of lipolysis and suppression of liver glucose output has been reported in dogs (Rebrin *et al.,* 1996), in rats (Lam *et al*., 2002; Lam *et al*., 2003) and in humans (Lewis *et al*., 1997).

In adult liver, expression of the enzyme lipoprotein lipase (Lpl) is physiologically relatively low (Liu *et al*., 2016) but can be increased by insulin (Ramos *et al*., 1999) or pro-inflammatory molecules such as TNFα (Enerbäck *et al*., 1988). We found that *Lpl* gene expression is up-regulated in liver of P2Y6^ΔNcr1^ mice in the clamped (insulin-stimulated) state but not in basal (unstimulated) conditions (data not shown). Furthermore, it is known, that the activity of Lpl is modulated by apolipoproteins such as apolipoprotein A-V (Apoa5) (Fruchart-Najib *et al*., 2004) whose expression in turn is regulated by insulin via phosphoinositide 3-kinases (PI3K) (Nowak *et al*., 2005) or on a transcriptional level by the nuclear receptor RAR Related Orphan Receptor alpha (Rorα) (Genoux *et al*., 2005). In our transcriptomic studies of clamped liver, we found that the expression of both these genes *Apoa5* and *Rora* is down-regulated in P2Y6^ΔNcr1^ mice. These observations further support the finding that hepatic insulin sensitivity is improved in these mice, while insulin-stimulated AKT activation remained unaltered.

Collectively, our data show that depletion of P2Y6R in NK cells reduces metaflammation in adipose tissue of HFD-fed mice leading to an overall improvement in insulin sensitivity. This improvement in insulin sensitivity is mainly reflected by a reduction of lipolysis in PGAT, an improved insulin-mediated suppression of HGP and the regulation of insulin-dependent genes, despite the notion that assessment of insulin-stimulated AKT activation upon injection of a high dose of insulin was comparable between the mice of the different genotypes.

NK cells are a heterogeneous innate lymphocyte cell population with organ- and microenvironment-specific expression patterns of cytotoxic molecules, receptors and transcription factors (Crinier *et al*., 2018; Zhao *et al*., 2020). Despite this tissue-specificity, we found a common set of ten genes that were similarly differentially expressed in adipose tissue and liver of clamped, thus insulin-stimulated mice. As we assumed that this gene set reflects that of tissue-infiltrating NK cells, rather than being an insulin-regulated gene set, we further investigated the transcriptome of isolated NK cells from adipose tissue and liver of HFD-fed P2Y6^fl/fl^ and P2Y6^ΔNcr1^ mice. Also here, transcripts of this gene set were found to be differentially expressed in NK cells of both origins, supporting our hypothesis that this gene set is NK cell-specific and that the deletion of P2Y6R from NK cells might alter the intrinsic expression pattern of tissue infiltrating NK cells. There are multiple unique lineages of NK cells, e.g. circulating conventional (c)NK cells, thymic NK cells, tissue-resident (tr)NK cells of the liver and skin and uterine (u)NK cells (Sojka, Tian, Yokoyama, 2014; Peng and Tian, 2017; Hashemi and Malarkannan, 2020). Each of these NK cell populations is characterized by a unique repertoire of receptors and cytokines and seems to emerge from a distinct developmental pathway (Erick and Brossay, 2016). Conventional NK cells are characterized by the expression of DX5 (CD49b) and are found as circulating cells in the blood or as tissue-infiltrating cells in different organs (Sojka *et al*., 2014; Erick and Brossay, 2016). In contrast to that, tissue-resident NK cells are characterized by the expression of CD49a and exhibit a rather immature phenotype (Sojka, Tian, Yokoyama, 2014; Sojka *et al*., 2014). Despite a large amount of studies with RNASeq data of NK cells from different origins such as blood (Yang *et al*., 2019; Smith *et al*., 2020; Zhao *et al*., 2020), liver (Filipovic *et al*., 2019; Zhao *et al*., 2020) and spleen (Peng *et al*., 2013; Sojka *et al*., 2014; Zhao *et al*., 2020), as well as from tumor infiltrating NK cells (de Andrade *et al*., 2019; Cursons *et al*., 2019), we could assign none of the ten genes of our NK cell-specific gene cluster to any of these available gene sets with the exception of *Rora*, whose expression has been reported to be higher in liver-resident DX5^-^ NK cells (Peng *et al*., 2013).

The only gene that was consistently up-regulated in all our transcriptomic data sets was *Lilra5.* This receptor belongs to the family of leukocyte immunoglobin-like receptors which are known to regulate leukocyte activation. Lilra5 has been reported to exist as transmembrane protein or as secreted molecule (Borges *et al*., 2003). The ligands for some but not all LILRs are classical and non-classical HLA-class I molecules, however the ligand for Lilra5 is still unknown (Kaur *et al*., 2013). Cross-linking Lilra5 to the surface of macrophages has been reported to trigger an increase in intracellular calcium concentrations and the release of cytotoxic molecules such as TNFα and IL-10 (Mitchell *et al*., 2008). Our data shows clear expression of *Lilra5* in P2Y6R-deficient NK cells, however in literature it is reported to be mainly expressed in macrophages and to some extend in B cells and granulocytes (Human Protein Atlas, https://www.proteinatlas.org/) and functional studies on Lilra5 until now are limited.

Collectively, our findings underline the importance of further studies to unravel the role of P2Y6R-signaling in NK cells especially in regard to the regulation of this here reported NK cell-specific gene cluster. In light of the finding that cell numbers in spleen and liver of P2Y6^ΔNcr1^ mice are reduced, further studies on conventional and tissue-resident NK cells and their infiltration kinetics in dependence of P2Y6R-signaling should be conducted to further define the impact of NK cell-specific P2YR-signaling on obesity-induced metaflammation.

Several publications have reported a role for a P2Y6R-dependent regulation of cytokine expression and release in macrophages (Bar *et al*., 2008; Lattin *et al*., 2008). In light of macrophages’ roles as drivers for obesity-induced metaflammation (Xu *et al*., 2003) and an increase in UDP levels in adipose tissue upon HFD-feeding, it was an obvious question whether macrophage-specific P2Y6R-signaling might contribute to obesity-associated metaflammation and development of insulin resistance. However, here we showed that mice with macrophage-specific abrogation of P2Y6R-signaling did not present in any conspicuous phenotype in regard to bodyweight development, insulin sensitivity, food intake or energy expenditure. In addition, we did not observe any alterations in adipose tissue inflammation. As obesity development is accompanied by a macrophage switch from M2- to M1-polarized macrophages in adipose tissue (Castoldi *et al*., 2016), it was interesting to note that the expression of P2Y6R is 3-fold up-regulated in M2-polarized macrophages compared to unpolarized or M1-polarized macrophages *in vitro*. Comparison of M1 and M2 polarized macrophages derived from human blood only reported a difference in the expression of other P2Y receptors such as P2Y1R, P2Y12R and P2Y13R that were found to be down-regulated in M1 compared to M2 macrophages and P2Y2R, P2Y8R, P2Y10R and P2Y14R being up-regulated (Gerrick *et al*., 2018). Another study on murine BMDMs found P2Y1R to be up-regulated in M2-polarized macrophages when compared to unpolarized cells (Jablonski *et al*., 2015). Nevertheless, our data clearly indicate that P2Y6R signaling in macrophages appears to be dispensable for the manifestation of obesity-induced metaflammation.

Taken together, our study demonstrates that insulin sensitivity and metaflammation depend on P2Y6R signaling in NK cells in the context of obesity, which may provide a target for the treatment of obesity-associated pathologies and diabetes.

## Materials and Methods

### Animals

All mouse experiments were approved by the local authorities (Bezirksregierung Köln, Germany) and conducted in accordance with NIH guidelines. Mice were housed in groups of 3-5 animals at 22-24°C and a 12-hour light/dark cycle. Mice had *ad libitum* access to food and water at all times, and food was only withdrawn if required for an experiment. Mouse models were generated by crossing previously published mouse strains, all of which have been backcrossed to C57BL/6 mice for at least ten generations. Briefly, mice with a conditional knockout of the P2Y6R gene in NK cells (P2Y6^ΔNcr1^) were generated by crossing Ncr1-Cre mice (Eckelhardt *et al*., 2011) with P2Y6R-floxed mice (Jain *et al*., 2020) and line breeding was maintained with Ncr1-Cre^+/-^:P2Y6^fl/fl^ crossed with Ncr1-Cre^-/-^:P2Y6^fl/fl^ mice. Mice with a conditional knockout of the P2Y6R gene in macrophages (P2Y6^ΔLysM^) were generated by crossing LysM-Cre mice (Clausen *et al*., 1999) with P2Y6R-floxed mice (Jain *et al*., 2020) and line breeding was maintained with LysM-Cre^+/-^:P2Y6^fl/fl^ crossed with LysM-Cre^-/-^:P2Y6^fl/fl^ mice. Genotyping of mouse lines was performed by genomic PCR on DNA isolated from tail biopsies using the primers listed in table 1. C57BL/6 N mice were ordered from Charles River (Cologne, Germany).

**Table 1:**
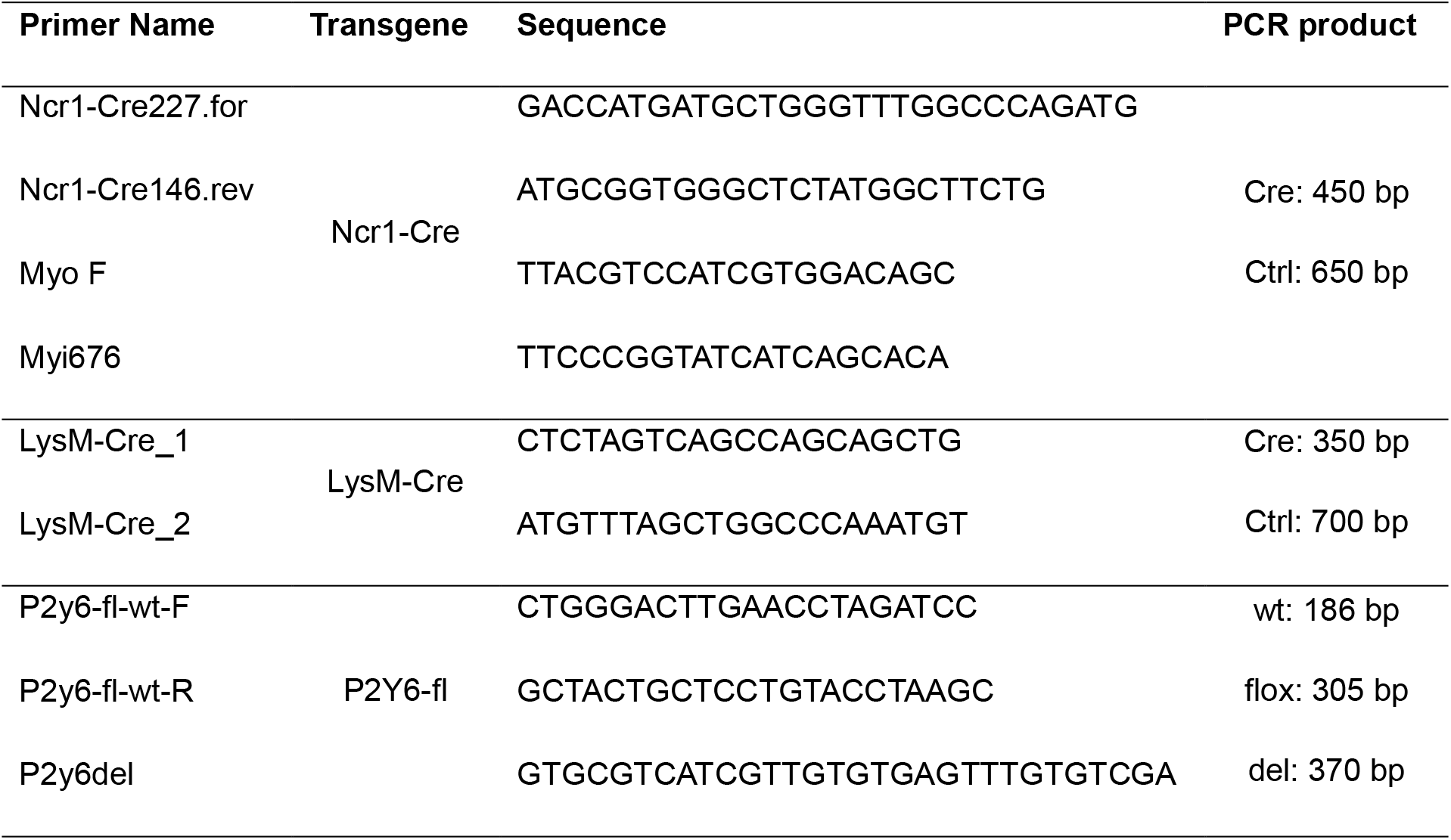
Genotyping primers.

### Metabolic Phenotyping

#### Diet Induced Obesity and High Fat Diet Feeding

Mice were fed a normal chow diet (NCD) (ssniff R/M-H Phytoestrogenarm, ssniff Spezialdiäten GmbH, Soest, Germany) until used for experiments. Experimental animals were either fed a high-fat diet (HFD) (ssniff D12492 (I) mod., ssniff Spezialdiäten GmbH, Soest, Germany) or a low-fat, low-sugar control diet (CD) (ssniff D12450B mod. LS, ssniff Spezialdiäten GmbH, Soest, Germany) starting at the age of 6 weeks.

#### Body Composition Analysis

Body weights were assessed weekly. Body composition was analyzed by computed tomography (CT) in isoflurane-anesthetized mice (Dräger Healthcare, Cologne, Germany). For data acquisition on an IVIS Spectrum CT scanner (Caliper LifeScience, Hopkinton, MA, USA) we used IVIS LivingImage Software V4.3.1. Quantification of lean and fat mass contents were determined with a modification of the previously described Vinci software package 5.06.0 (Cízek *et al*., 2004) developed at our institution (available at: www.nf.mpg.de/vinci3).

#### Glucose and Insulin Tolerance Test, and HOMA-IR Determination

Glucose tolerance tests were performed in 16 hours overnight-fasted mice after 9, 14 and 20 weeks of HFD- or CD-feeding. Fasted blood glucose concentrations (mg/dl) were determined prior to the *i.p.* injection of 20% (w/v) glucose (B.Braun, Melsungen, Germany; 2 mg/g body weight). Fifteen, 30, 60 and 120 min after injection, blood glucose levels in tail vein blood were measured using an automatic glucometer (Bayer Contour Next, Bayer AG, Leverkusen, Germany). Insulin tolerance tests were performed in random fed mice at corresponding ages as GTTs. Blood glucose concentrations (mg/dl) were determined prior to the test. Before *i.p.* injection of insulin (0.5 mU/g body weight; Insuman Rapid 40 I.E./ml, Sanofi, Paris, France), food was removed from the cages for the duration of the experiment. Blood glucose levels were measured 15, 30 and 60 min after injection. HOMA-IR is a method to quantify insulin resistance. Here, HOMA-IR was calculated using the formula (Insulin_fasted_ [µU/ml] x Glucose_fasted_ [mg/dl]) x 405.

#### Hyperinsulinemic-Euglycemic Clamp Analyses

Surgical implantation of catheters in the jugular vein was performed as previously described (Könner *et al*., 2007). After one week of recovery, mice were fasted for 4 hours in the morning of the day of experiment, which was conducted based on the protocol from the Mouse Metabolomic Phenotyping Center at Vanderbilt University (Ayala *et al*., 2011). Prior to the actual clamp, mice received 270 µl of a 3% (v/v) plasma solution at a rate of 0.18 ml/h for 90 min in parallel with a primed-continuous tracer D-[3-^3^H]-Glucose infusion (PerkinElmer, Waltham, MA, USA, 0.8 μCi prime, 0.04 μCi/min). After this basal period, a blood sample was collected from the tail tip for determination of basal parameters. Following, the clamp started with the infusion of an insulin solution (Sanofi, Paris, France or Lilly, Bad Homburg, Germany; 6 μU/g/min) together with a 40% glucose solution (Bela Pharm, Vechta, Germany) primed with D-[3-^3^H]-Glucose (0.06 ml/h). After 30 min of infusion, venous blood glucose levels were determined every 10 min using an ultra-sensitive glucometer (B-Glucose Analyzer; Hemocue, Ängelhom, Sweden) and glucose infusion rates were adjusted in order to keep blood glucose at physiological levels (±140 mg/dl). The steady state was considered, when a fixed glucose infusion rate maintained the blood glucose levels constant (± 140 mg/dl) for 30 min. Approximately 90 min after start of the insulin infusion, steady state levels were reached. During the steady state, blood samples were collected every 10 min for the measurement of steady-state parameters. Then, animals received a 100 µl bolus of 2-[1-^14^C]-Deoxy-D-glucose (American Radiolabeled Chemicals Inc., St. Louis, MO, USA; 10 μCi) at a rate of 1.2 ml/h. Following, infusion was switched back to insulin (6 µU/g/min) and 40% glucose primed with D-[3-^3^H]-Glucose (40 µl bolus at 1.2 ml/h followed by continuous infusion at the rate used to keep steady state glucose levels). Blood samples were collected and blood glucose levels determined at 0, 5, 10, 15, 25 and 35 min after this last bolus for follow-up determination of 2-[1-^14^C]-Deoxy-D-glucose withdrawal from plasma.

At the end of the experiment, mice were sacrificed by cervical dislocation, and liver, perigonodal adipose tissue, brain, brown adipose tissue and skeletal muscle were dissected and stored at −80°C. Plasma [3-^3^H]-Glucose radioactivity of basal and steady state was measured as described by the Vanderbilt protocol (Ayala *et al*., 2011). Plasma 2-[1-^14^C]-Deoxy-D-glucose radioactivity was directly measured in a liquid scintillation counter (Tri-Carb 2810 TR, Perkin Elmer, Waltham, MA, USA). Tissue lysates were processed through ion exchange chromatography columns (Bio Rad Laboratories Inc., Hercules, CA, USA) to separate 2-[1-^14^C]-Deoxy-D-glucose (2DG) from 2-[1-^14^C]-Deoxy-D-glucose-6-phosphate (2DG6P). Glucose infusion rate (GIR) and hepatic glucose production (HGP) was calculated as previously described (Könner *et al*., 2007). *In vivo* glucose uptake into perigonodal and brown adipose tissue, brain, liver and skeletal muscle was calculated based on the accumulation of 2DG6P in the respective tissue and the disappearance rate of 2DG from plasma as described previously (Könner *et al*., 2007).

#### Analyses of Plasma Samples

Blood was collected in EDTA-treated tubes (Sarstedt, Nümbrecht, Germany) and centrifuged for 10 min at 17 000 g and 4°C. The plasma was immediately frozen in liquid nitrogen and stored at −80°C until used for ELISA analysis. Fasted mouse insulin and leptin plasma concentrations were determined by ELISA with mouse standards according to the manufactureŕs guidelines (Ultra Sensitive Mouse Insulin ELISA Kit, Crystal Chem, Elk Grove Village, IL, USA; Mouse Leptin ELISA Kit, Crystal Chem, Elk Grove Village, IL, USA). For the leptin ELISA, plasma samples were diluted 1:20, for the insulin ELISA undiluted plasma samples were used. The concentration of human insulin in plasma of clamped mice was determined by ELISA according to manufactureŕs guidelines (Insulin (Human) Ultrasensitive, DRG Diagnostics, Marburg, Germany) using 1:25 diluted plasma samples. Free fatty acid concentration in plasma (diluted 1:25) of clamped mice was determined using a quantitative Free Fatty Acid Assay Kit (Abcam, Cambridge, Great Britain). Absorbance was measured with the Filter Max F5 device (Molecular Devices, San José, CA, USA) at corresponding wavelengths. Concentrations were calculated using a 4-parameter logistic regression analysis.

#### Indirect Calorimetry

Indirect calorimetry was performed using an open-circuit, indirect calorimetry system (PhenoMaster, TSE systems, Bad Homburg vor der Höhe, Germany). Three days before data acquisition, mice were adapted to the food/drink dispensers of the PhenoMaster system and the body weight was monitored. Then, mice were placed into regular type II cages with sealed lids and allowed to adapt to the chambers at least 24 hours at room temperature (22°C). During data acquisition, mice were provided with food and drink *ad libitum*. All parameters were measured simultaneously and continuously.

### Gene Expression Analyses

#### RNA Extraction from Tissues and Cells

Snap frozen tissue samples or cells were homogenized in 0.5-1 ml Qiazol (Qiagen, Hilden, Germany) using a Retsch Homogenizer (Retsch GmbH, Haan, Germany) combined with beads (Bertin Technologies, Montigny-le-Bretonneux, France). Afterwards, 0.2-0.4 ml chloroform were added to each sample and the mixture incubated for 5 min at RT. Following, samples were centrifuged for 15 min at 4°C and 12 200 g. The supernatant was transferred to a new tube and mixed with the equal volume of 70% ethanol. The total volume was then transferred to a RNeasy Mini Spin Column of the RNeasy Mini Kit (Qiagen, Hilden, Germany). Subsequent steps were conducted as described in the manufactureŕs protocol. DNAse treatment (15 min at RT; Qiagen, Hilden, Germany) was only conducted in samples used for RNASeq but not in samples used for qPCR analysis. Finally, RNA was eluted in H_2_O and stored at −80°C until further use.

#### cDNA Synthesis and Quantitative Real Time PCR

Concentrations of RNA extracted from PGAT, liver or BMDMs were determined with the NanoDrop ND-1000 device (Thermo Fisher Scientific, Waltham, MA, USA). cDNA synthesis was conducted using the High Capacity cDNA RT kit according to manufactureŕs guidelines (Applied Biosystems, Foster City, CA, USA). Following, cDNA was used for real-time PCR analysis using Takyon Low Rox Probe 2x Master Mix dTTP Blue (Eurogentec, Liège, Belgium) with TaqMan® Assay-on-demand kits (Applied Biosystems, Foster City, CA, USA). Relative expression of target mRNAs was adjusted for total RNA content by *hypoxanthine guanine phosphoribosyl transferase 1* (*Hprt1*) or *TATA binding protein* (*Tbp*) mRNA qRT-PCR (for target probe details see table 2). Calculations were performed by a comparative method (2^−ΔΔCT^). Quantitative PCR was performed on a QuantStudio 7 Flex Real-Time PCR System using the QuantStudio Real-Time PCR Software v1.3 (Life Technologies, Carlsbad, CA, USA).

**Table 2:** Probes used for quantitative real time PCR.

| Target gene | Probe ID | Fluorophore | Company |
| --- | --- | --- | --- |
| *P2y6r* | Mm02620937_s1 | FAM | Thermo Fisher Scientific |
| *Il6* | Mm00446190_m1 | FAM | Thermo Fisher Scientific |
| *Tnfa* | Mm00443258_m1 | FAM | Thermo Fisher Scientific |
| *Ccl2* | Mm00441242_m1 | FAM | Thermo Fisher Scientific |
| *Ccl3* | Mm00441258_m1 | FAM | Thermo Fisher Scientific |
| *Ifng* | Mm01168134_m1 | FAM | Thermo Fisher Scientific |
| *Il10* | Mm00439614_m1 | FAM | Thermo Fisher Scientific |
| *Il12a* | Mm00434165_m1 | FAM | Thermo Fisher Scientific |
| *Il1b* | Mm01336189_m1 | FAM | Thermo Fisher Scientific |
| *Lpl* | Mm00434764_m1 | FAM | Thermo Fisher Scientific |
| *Rora* | Mm01173766_m1 | FAM | Thermo Fisher Scientific |
| *Plin2* | Mm00475794_m1 | FAM | Thermo Fisher Scientific |
| *Lilra5* | Mm01335361_m1 | FAM | Thermo Fisher Scientific |
| *Hprt* | Mm01545399_m1 | FAM | Thermo Fisher Scientific |
| *Hprt* | Mm03024075_m1 | VIC | Thermo Fisher Scientific |
| *Tbp* | Mm00446973_m1 | VIC | Thermo Fisher Scientific |

#### RNA Sequencing of Clamped Tissue Samples

RNA from clamped liver and PGAT samples was isolated as described above. Libraries were prepared using the TruSeq Stranded mRNA protocol according to the manufactureŕs recommendations (Illumina, San Diego, CA, USA). After validation with the Agilent 4200 TapeStation (Agilent, Santa Clara, CA, USA) and quantification with a Qubit device (Invitrogen, Carlsbad, CA, USA), all transcriptome libraries were pooled. The pool was quantified using the Peqlab KAPA Library Quantification Kit (VWR, Radnor, PA; USA) and the Applied Biosystems 7900HT Sequence Detection (ThermoFisher, Waltham, MA, USA) and sequenced on a Illumina NovaSeq S2 flowcell (Illumina, San Diego, CA, USA) with a PE100 protocol.

### Immunoblots

#### Protein Extraction from Tissue Samples

Snap frozen tissue samples were homogenized in 1 ml lysis buffer containing 50 mM Tris, 130 mM NaCl, 5 mM EDTA, 1% (v/v) NP-40, 1 mM NaF, 1 mM PMSF (Sigma Aldrich, St. Louis, MO, USA), cOmplete mini protease inhibitor cocktail (1 tablet/10 ml; Roche, Basel, Switzerland) and PhosSTOP Phosphatase Inhibitor Cocktail (1 tablet/10 ml; Roche, Basel, Switzerland) using a Retsch Homogenizer (Retsch GmbH, Haan, Germany) combined with ceramic beads (Bertin Technologies, Montigny-le-Bretonneux, France). After homogenization for 2 min at a frequency of 30 1/s, homogenates were incubated for 20 min on ice followed by centrifugation at 13 000 rpm for 20 min at 4°C. Protein concentration was determined relative to a BSA standard using the Pierce^TM^ BCA^TM^ Protein-Assay (Thermo Scientific, Waltham, MA, USA). Lysates were finally adjusted to a final concentration of 2 µg/µl with 4X Laemmli buffer (Bio-Rad, Hercules, CA, USA) containing 10% β-Mercaptoethanol.

#### Western Blotting

Proteins (20 µg per well) were separated in 4–15% Criterion™ TGX™ Precast Midi Protein Gels with 26 wells (Bio-Rad, Hercules, CA, USA) at a constant voltage of 120 V. The SDS-Running buffer contained 200 mM Glycine, 25 mM Tris and 1% (w/v) SDS. Proteins were transferred to Trans-Blot Turbo Midi 0.2 µm PVDF Transfer membranes (Bio-Rad, Hercules, CA, USA) using the Trans-Blot Turbosystem (Bio-Rad, Hercules, CA, USA). Following, membranes were blocked in western blotting reagent (Roche, Basel, Switzerland) diluted 1:10 in TBS-T buffer (Tris-buffered saline with 1% (v/v) Tween 20) for 1–2 h at room temperature. Incubation with primary antibodies was conducted over night at 4°C. Subsequently, membranes were washed three times for 5–7 min in TBS-T followed by the incubation in horse radish peroxidase-labeled secondary antibody over night at 4°C. All primary and secondary antibodies were diluted in 5% (v/v) western blotting reagent (Roche, Basel, Switzerland). After washing for 3 times in TBS-T, membranes were incubated with SuperSignal ECL Western Blotting Substrate (Thermo Fisher Scientific, Waltham, MA, USA) and luminescence was detected by the Fusion Solo Vilber Lourmat system (Vilber Lourmat GmbH, Eberhardzell, Germany). If necessary, membranes were stripped in stripping buffer (6% SDS (v/v), 188 mM Tris pH 6.8, 2% (v/v) β-Mercaptoethanol) for 30 min at 56°C in a shaking water bath. Afterwards, membranes were washed three times in TBS-T followed by blocking in 10% (v/v) western blotting reagent (Roche, Basel, Switzerland) for 1–2 h at room temperature. Then, the incubation with primary and secondary antibodies was conducted as described above. Band intensities were quantified using ImageJ (NIH). The intensity of protein bands was normalized to the intensity of calnexin or β-Actin bands which served as loading control. Band intensities of protein bands from phospho-targets such as pAKT and pHSL were additionally normalized to bands of total protein, AKT and HSL, respectively. For detailed information of individual primary and secondary antibodies see table 3.

**Table 3:**
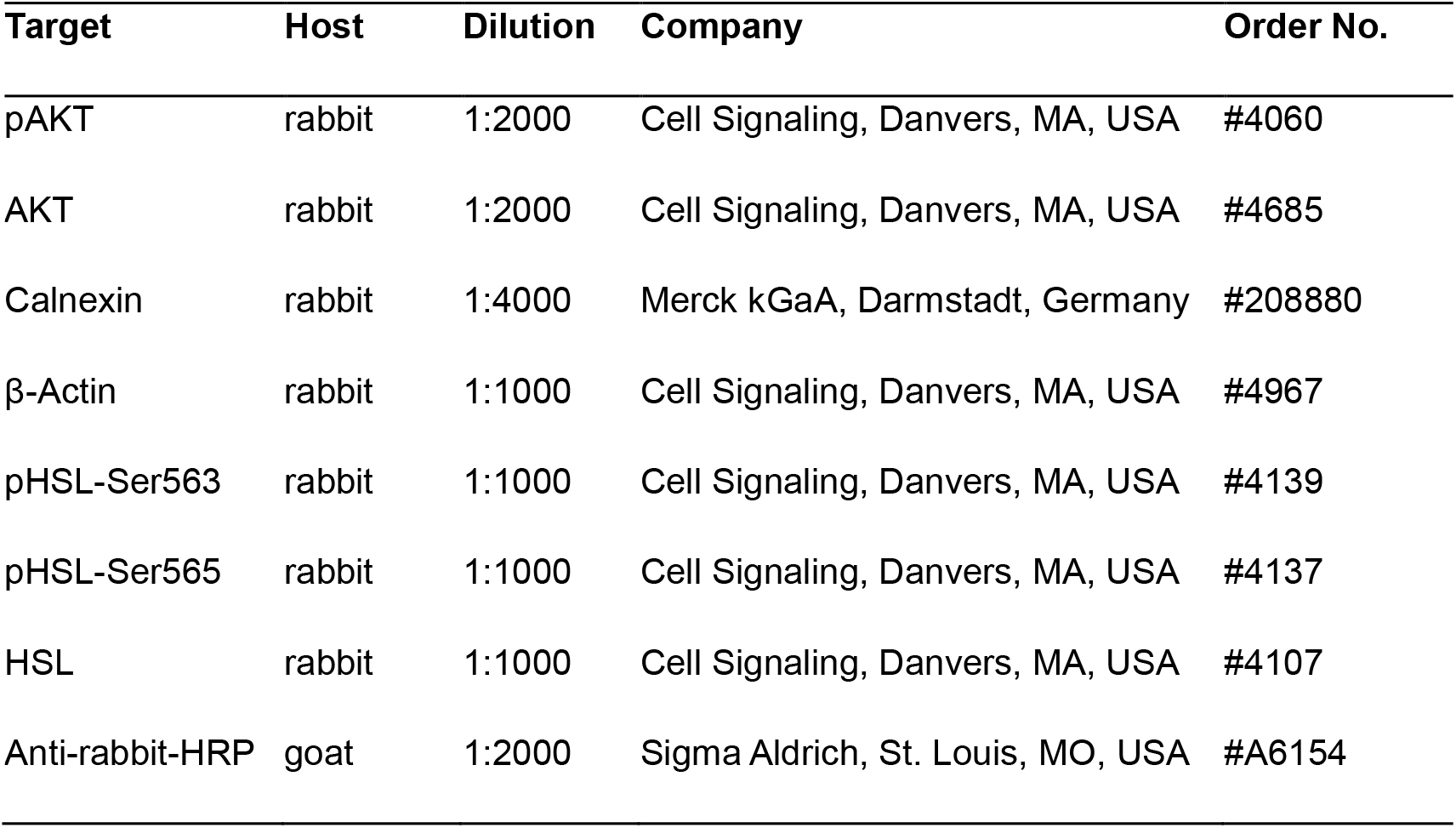
Detailed description of antibodies used for Western Blots.

### Analyses of Isolated Immune

#### Cells Immune Cell Isolation from Tissue

Leucocytes were isolated from perigonodal adipose tissue (PGAT), liver, spleen and blood from HFD-fed male mice according to the protocol described in Theurich *et al*., 2017. Briefly, liver and PGAT were dissociated in lysis buffer (liver: 150 mM NaCl, 5.6 mM KCl, 5.5 mM Glucose, 20.1 mM HEPES, 25 mM NaHCO_3_, 2 mM CaCl_2_, 2 mM MgCl_2_, 500 U/ml Collagenase IV; PGAT: 120 mM NaCl, 5.2 mM KH_2_PO_2_, 1 mM MgSO_4_, 10 mM Hepes, 10 mM NaHCO_3_, 500U/ml Collagenase I) with a tissue-lyser (Gentle MACS, Miltenyi Biotec, Bergisch Gladbach, Germany) and digested enzymatically for 20-30 min at 37°C while continuously shaking in a water bath. The spleen was dissociated by pressing it through a 100 µm nylon mesh (Greiner Bio-One, Solingen, Germany) and cells were collected in FACS buffer (autoMACS Running Buffer, Miltenyi Biotec, Bergisch Gladbach, Germany). Blood was collected in EDTA-treated tubes (Sarstedt, Nümbrecht, Germany). Immune cells were separated by centrifugation: adipose tissue homogenates at 400 g and 18°C for 5 min, liver homogenates via 20% (w/v) Histodenz (Sigma Aldrich, St. Louis, MO, USA) density-centrifugation at 1500 g and 4°C, spleen homogenates at 400 g and 4°C for 5 min. Following, cells and blood were subjected to erythrocyte lysis in RBC lysis buffer (BioLegend, San Diego, CA, USA) for 10 min on ice. Erythrocyte lysis was terminated by washing the cells in the 10-fold volume of FACS buffer (autoMACS Running Buffer, Miltenyi Biotec, Bergisch Gladbach, Germany) and passed through a 30 µm cell strainer (BD Biosciences, Franklin Lakes, USA) to remove large cellular debris. After centrifugation at 400 g and 4°C for 5 min, cells were resuspended in 1 ml FACS buffer (autoMACS Running Buffer, Miltenyi Biotec, Bergisch Gladbach, Germany) and used for cell counting.

#### Flow Cytometry

For flow cytometry analysis, 1×10^5^-1×10^6^ cells were stained after Fc-blocking with TruStain FcX (BioLegend, San Diego, CA, USA) and incubation with Zombie Aqua fixable viability dye (BioLegend, San Diego, CA, USA). Fluorochrome-conjugated antibodies were used for specific immunostainings with dilutions (in autoMACS Running Buffer, Miltenyi Biotec, Bergisch Gladbach, Germany) as listed in table 4: anti-mCD45 (30-F11), anti-mCD3 (17A2), anti-mCD19 (6D5), anti-mNK1.1 (PK136), anti-mNcr1 (29A1.4), anti-mCD11b (M1/70), anti-mCD27 (LG.3A10) and anti F4/80 (BM8). Cells were analyzed using an 8-color flow cytometer (MACSQuant10, Miltenyi Biotec, Bergisch Gladbach, Germany) and data analysis was performed with FlowJo 10.4 (FlowJo LLC, BD Biosciences, San Jose, CA, USA) software packages.

**Table 4:** Detailed description of antibodies used for flow cytometry.

| Target | Fluorophor | Dilution | Clone | Company | Order No. |
| --- | --- | --- | --- | --- | --- |
| NK1.1 | BV421 | 1:50 | PK136 | BioLegend, San Diego, CA, USA | #108732 |
| CD19 | PE/Cy7 | 1:50 | 6D5 | BioLegend, San Diego, CA, USA | #115520 |
| CD3 | APC | 1:100 | 17A2 | BioLegend, San Diego, CA, USA | #100236 |
| CD45 | APC/Cy7 | 1:100 | 30-F11 | BioLegend, San Diego, CA, USA | #103115 |
| CD11b | PerCP/Cy5.5 | 1:100 | M1/70 | BioLegend, San Diego, CA, USA | #101227 |
| CD27 | PE/Cy7 | 1:50 | LG.3A10 | BioLegend, San Diego, CA, USA | #124215 |
| Live/Dead | ZombieAqua | 1:100 | - | BioLegend, San Diego, CA, USA | #423102 |
| F4/80 | PE | 1:50 | BM8 | BioLegend, San Diego, CA, USA | #123110 |
| Ncr1 | BV421 | 1:20 | 29A1.4 | BioLegend, San Diego, CA, USA | #137611 |
| NK1.1 | Alexa488 | 1:50 | PK136 | BioLegend, San Diego, CA, USA | #108718 |
| TruStain FcX | - | 1:100 | - | BioLegend, San Diego, CA, USA | #101320 |
| CD3 | PerCP/Cy5.5 | 1:100 | 17A2 | BioLegend, San Diego, CA, USA | #100218 |
| CD11b | APC | 1:100 | M1/70 | BioLegend, San Diego, CA, USA | #101212 |

#### RNA Isolation and RNASeq of Bulk Sorted NK Cells

NK cells from murine organs and blood were purified from single cell suspensions using FACSAria IIIu or FACSAria Fusion cell sorters (BD Biosciences, San Jose, CA, USA) after immunostainings as described above. NK cells were sorted from HFD-fed animals identified as single/viable/CD45^+^/CD3^-^/NK1.1^+^/Ncr1^+^ cells using a 70 µm nozzle at 70 psi pressure. During sorting, sample and collection tubes were chilled. Purified cells were directly sorted into FACS buffer (autoMACS Running Buffer, Miltenyi Biotec, Bergisch Gladbach, Germany) and pelleted by centrifugation for 5 min at 400 g and 4°C. Total RNA was extracted using the Arcturus RNA picopure kit (Thermo Fisher Scientific, Waltham, MA, USA) following the manufacturer’s instructions. Due to the low amount of input material, pre-amplification using the Ovation RNASeq System V2 (NuGEN, Redwood, CA, USA), was performed as described previously (Paeger *et al*., 2017). For library preparation, the Illumina Nextera XT DNA sample preparation protocol (Illumina, San Diego, CA, USA) was used, with 1 ng cDNA input. After validation with the Agilent 4200 TapeStation (Agilent, Santa Clara, CA, USA) and quantification with a Qubit device (Invitrogen, Carlsbad, CA, USA) all libraries were pooled. The pool was quantified using the Peqlab KAPA Library Quantification Kit (VWR, Radnor, PA; USA) and the Applied Biosystems 7900HT Sequence Detection (ThermoFisher, Waltham, MA, USA) and sequenced on an Illumina NovaSeq system (Illumina, San Diego, CA, USA) with PE100 read length.

#### Analysis of RNA-Sequencing Results

We applied the community-curated nfcore/rnaseq analysis pipeline version 1.4.2 (Ewels *et al*., 2019) for processing RNA-sequencing data. The gene-level quantification was carried out using Salmon 0.14.1 (Patro *et al*., 2017) using the reference genome GRCm38. The differential gene expression analysis was done using the DESeq2 1.26.0 (Love *et al*., 2014) R package. We denote a gene differentially expressed with an FDR adjusted_p_value≤0.05. Gene-ontology term analysis of the differentially expressed genes was carried out using the clusterProfiler 3.14.3 (Yu *et al*., 2012) R package.

#### NK Cell Isolation by Magnetic Separation and DNA Extraction

For knock out validation, NK cells were isolated from spleen of P2Y6^fl/fl^ and P2Y6^ΔNcr1^ mice fed a normal chow diet using the mouse NK cell isolation kit from Miltenyi Biotec, (Bergisch Gladbach, Germany). The spleen was dissociated by pressing it through a 100 µm nylon mesh (Greiner Bio-One, Solingen, Germany). Following, erythrocyte lysis was carried out for 10 min on ice in RBC lysis buffer (BioLegend, San Diego, CA, USA). Cells were centrifuged for 5 min at 400 g and 4°C. The pellet was resuspended in FACS buffer (autoMACs Running Buffer, Miltenyi Biotec, Bergisch Gladbach, Germany) and cells were passed through a 30 µm nylon mesh to remove cell clumps. After a second centrifugation at 400 g for 5 min at 4°C, cells were resuspended in 1 ml FACS buffer (autoMACS Running Buffer, Miltenyi Biotec, Bergisch Gladbach, Germany) and used for cell counting. Following, NK cells were isolated according to the manufactureŕs protocol (Miltenyi Biotec, Bergisch Gladbach, Germany). As 3 x 10^7^ cells were used per sample, all reagent volumes and total volumes were scaled up accordingly. Magnetic separation was carried out using LS columns (Miltenyi Biotec, Bergisch Gladbach, Germany). After separation, DNA was extracted from isolated NK cells and cells of the flow through (nonNK cells). Cells were lysed in lysis buffer (100 mM Tris-HCl pH 8.5, 5 mM EDTA pH 8.0, 0.2% SDS (v/v), 200 mM NaCl) at 37°C over night. DNA was precipitated in 0.5 ml ice-cold isopropanol followed by incubation in 0.5 ml 70% ethanol. The DNA pellet was resuspended in H_2_O and used for genomic PCR with primers listed in table 1.

#### Seahorse Analyses of Liver Micro-Punches

To obtain consistent liver punches for analysis, we used a 0.5 mm EMS-Core Sampling Tool (Electron Microscopy Science, Hatfield, PA, USA). Four to five tissue punches per animal were carefully moved into Seahorse XF96e spheroid microplates (Agilent, Santa Clara, CA, USA) containing Seahorse base medium (Agilent, Santa Clara, CA, USA) with 25 mM glucose, 1 mM sodium pyruvate and 2 mM L-glutamine. The spheroid microplates (Agilent, Santa Clara, CA, USA) were coated with Cell-Tak (22.4 mg/mL; Corning, Corning, NY, USA) to prevent detachment of the tissue punches during the assay. Plates were then transferred to a CO_2_-free incubator and maintained at 37°C for 30 min before starting the assay. Following instrument calibration, plates were transferred to the Seahorse XF96 Flux Analyzer (Agilent, Santa Clara, CA, USA) to record cellular oxygen consumption rates (OCR) and extracellular acidification rates (ECAR). Ten measurements were performed to assess baseline levels.

### Histological Analysis of PGAT and Liver

#### MAC2 Staining in PGAT

Perigonodal adipose tissue (PGAT) was isolated and fixed in 4% (w/v) paraformaldehyde for 2-3 days at 4°C. Fixed tissue samples were then embedded in paraffin and 5 µm thick sections were cut at a rotary microtome (Leica, Wetzlar, Germany). After a 10 min incubation in PBS, endogenous peroxidases were blocked for 20 min with 0.3% (v/v) H_2_O_2_ (in PBS and 0.25% (v/v) TritonX). After washing with PBS, sections were blocked with 10% (v/v) Rotiblock (Carl Roth, Karlsruhe, Germany) (diluted in PBS and 0.25% (v/v) TritonX) for 80 min. Following, primary antibody incubation in rat-anti-MAC2 antibody (Biozol, Eching, Germany; 1:100 diluted in Signalstain, Cell Signaling, Danvers, MA, USA) was conducted overnight at 4°C. After washing with PBS and 0.25% (v/v) Triton X, secondary antibody incubation with HRP-conjugated goat-anti-rat (Jackson Immuno Research, Cambridgeshire, UK; 1:100 diluted in PBS + 0.1% (v/v) TritonX) was carried out for 1h at room temperature. After washing in PBS and 0.25% Triton X followed by DAB reaction and staining of nuclei via Mayers Hematoxylin (Sigma Aldrich, St. Louis, MO, USA), sections were scanned on a Leica DM 1000 LED (Leica, Wetzlar, Germany).

#### Oil-Red-O Staining in Liver

After sacrifice, liver tissue was embedded in tissue freezing medium (Leica, Wetzlar, Germany) and frozen down on dry ice. Liver sections were cut on a cryostat (Leica, Wetzlar, Germany) as 7 µm thick sections. Prior to the staining, tissue sections were washed for 5 min in tap water, followed by an incubation in Oil-Red-O solution (0.3 g Oil-Red-O (Sigma Aldrich, St. Louis, MO, USA) in 60 ml isopropanol and 40 ml tap water) for 15 min. After several washes in tap water, sections were incubated for 5 min in Mayers Hematoxylin (Sigma Aldrich, St. Louis, MO, USA) followed by a 15 min incubation in tap water. Sections were embedded in Kaiseŕs glycerine gelatine (Merck Millipore, Burlington, MA, USA) and scanned on a Leica DM 1000 LED (Leica, Wetzlar, Germany).

#### Quantification of Immunohistochemical Stainings

MAC2 and Oil-Red-O stainings were quantified using Image J (NIH, Bethesda, MD, USA). First, images (RGB color) were split into different channels. Threshold was adjusted for the blue channel by finding a threshold value that fits at minimum three independent pictures. Integrated density values were used in order to compare genotypes. Each data point in Suppl. Fig.S8A and Suppl. Fig. S10 represents the mean integrated density value ± SEM of 2-3 technical replicates each of them originating from one biological replicate (one mouse).

The size of adipocytes was quantified using the Adiposoft plugin (Version 1.16; CIMA, University of Navarra) for Image J (NIH, Bethesda, MD, USA).

#### Bone Marrow-Derived Macrophage Cultures

Bone marrow was isolated from ethanol-rinsed (70% v/v) femurs and tibias of C57BL/6, P2Y6^ΔLysM^ or P2Y6^fl/fl^ mice fed a normal chow diet. Collected cells were centrifuged for 1 min at 300 g and 4°C, followed by red blood cell lysis in RBC lysis buffer (BioLegend, San Diego, CA, USA) for 5 min on ice. Following, cells were passed through a 30 µm cell strainer to remove cellular debris and pelleted by centrifugation at 1200 rpm for 5 min at 4°C. Then, cells were re-suspended in culture medium (RPMI 1640 w/o phenolred (Thermo Fisher Scientific, Waltham, MA, USA) supplemented with 10% FCS (v/v), 1% glutamine (v/v), 1% penicillin-streptomycin (v/v) and 50 ng/ml M-CSF (Miltenyi Biotec, Bergisch Gladbach, Germany) and seeded at a density of 5 x 10^5^ cells per well of a 6-well plate. Cells were differentiated for a minimum of 7 days with medium exchange every third day. One day before polarization experiments, BMDMs were washed twice with sterile PBS and further cultured in the absence of M-CSF. Cells were stimulated with 100 ng/ml LPS (Sigma Aldrich, St. Louis, MO, USA) and 20 ng/ml IFNγ (Thermo Fisher Scientific, Waltham, MA, USA) for M1-polarization or with 50 ng/ml IL-6 (R&D Systems, Minneapolis, MN, USA) and 20 ng/ml IL-4 (Miltenyi Biotec, Bergisch Gladbach, Germany) for M2-polarization. After 24 h of stimulation, cells were washed twice with sterile PBS and harvested using cell scarpers. Following centrifugation, cell pellets were stored at −80°C until further use.

#### Lipidomic Analyses of Liver Tissue

Chain-length specific DAG and TAG species from liver samples were quantified by nano-electrospray ionization tandem mass spectrometry (Nano-ESI-MS/MS) according to the protocol described in Turpin *et al*. 2019. Conditions of lipid extraction, Nano-ESI-MS/MS analysis and processing of mass spectra and quantification of lipid species was done as previously described (Kumar *et al*., 2015).

### Metabolomic Analyses

#### Sample collection

Mice were sacrificed after 6 and 12 weeks of HFD- or normal chow-diet feeding. A piece of liver tissue from the left lobe was dissected and directly snap frozen in liquid nitrogen. Perigonodal adipose tissue was dissected and dissociated in lysis buffer (120 mM NaCl, 5.2 mM KH_2_PO_2_, 1 mM MgSO_4_, 10 mM Hepes, 10 mM NaHCO_3_, 500U/ml Collagenase I) with a tissue-lyser (Gentle MACS, Miltenyi Biotec, Bergisch Gladbach, Germany, Germany) and digested enzymatically for 20-30 min at 37°C while continuously shaking in a water bath. Adipocytes and stromal vascular fraction cells were separated by centrifugation at 400 g at 18°C for 5 min. After centrifugation, the adipocytes accumulated as fat layer on the surface of the suspension and the SVF cells were pelleted at the bottom. Adipocytes were carefully transferred to a new vial an washed twice in PEB buffer (BSA 0.5% w/v, 2 mM EDTA, PBS) and then quick frozen in liquid nitrogen. The SVF cells were washed twice in PEB buffer and then snap frozen as pellet. Blood was collected in EDTA-treated tubes (Sarstedt, Nümbrecht, Germany) and plasma generated by centrifugation at 17 000 g for 10 min at 4°C, followed by snap freezing in liquid nitrogen. All samples were stored at −80°C until being used for metabolomic analyses.

#### Metabolite Extraction

Samples were aliquoted and, if necessary disintegrated with a ball mill (Qiagen, TissueLyser II). Each sample was extracted in 1000 µl of pre-cooled (−20°C) extraction buffer (UPLC-grade actonitrile:methanol:water [40:40:20 v/v]), which was spiked with 50 ng/ml of uridine d2 (Santa Cruz Biotechnology, sc-213138) and UDP-α-D-13C glucose (Cambridge Isotope Laboratories, CLM-10513) as internal standards. The sample was immediately vortexed until the plasma was homogenously suspended and samples were incubated for 30 min on an orbital mixer at 4°C and 1500 rpm, before sonicating them for 10 min in an ice-cooled bath-type sonicator. After sonication, samples were centrifuged for 10 min at 21.100 x g and 4°C and the cleared supernatant was transferred to fresh 1.5 mL Eppendorf tubes and concentrated to dryness using a Speed Vac concentrator (www.eppendorf.com). In parallel to the analytical samples, we prepared 50 µl aliquots of reference samples, which were extracted as described above. These reference samples contained distinct concentrations (0-5000 ng/ml) of the pure reference compounds uridine (Sigma Aldrich, St. Louis, MO, USA; U3750) and UDP (Sigma Aldrich, St. Louis, MO, USA; 94330).

#### Quantitative LC-MS Analysis of Uridine

For the mass spectrometric analysis of uridine levels, the concentrated metabolite pellet was re-suspended in 100 µl of UPLC grade water. Five µl of the cleared supernatant of the re-suspended samples was analyzed on an UPLC (Acquity iClass, Waters), using a HSST3 (100 x 1.0 mm column with 1.8 µm particle size, Waters) connected to a Xevo TQ-S (Waters) triple quadrupole mass spectrometer. The flow rate was set to 100 µl/min of buffer A (0.1% formic acid in UPLC grade water) and buffer B (0.1% formic acid in UPLC-grade acetonitrile) using the following gradient: 0-1 min 99-90% A; 1-5 min 90-50% A; 5-6.5 min 50-1%A; 6.6-7.5 min 1% A; 7.5-7.7 min 1-99% A. The column was re-equilibrated at 99% A for additional 3.3 min. The eluting metabolites were detected in positive ion mode using MRM (multi reaction monitoring) applying the following settings: capillary voltage 2.0 kV, desolvation temperature 550°C, desolvation gas flow rate 800 l/h, cone gas flow 150 ml/min, collision cell gas flow 0.15 ml/min. The following MRM transitions were used for relative compound quantification of uridine and uridine d2, respectively: m/z precursor mass (M+H^+^) 245/247, fragment mass (M+H^+^) m/z 113/115 using a cone voltage of 26/6 V and a collision energy of 16/10 V. For confirmation purpose of the analyzed compound an additional transition of the precursor mass (M+H^+^) m/z 245/247, fragment mass (M+H^+^) m/z 133/98 using a cone voltage of 26/6 V and a collision energy of 16/32 V was analyzed.

#### Anion-Exchange Chromatography-Mass Spectrometry (AEC-MS) Analysis of UDP

Parallel to the analysis of uridine an aqueous aliquot of the re-suspended metabolite pellet was analyzed using a Dionex ion chromatography system (ICS 5000+, Thermo Fisher Scientific, Waltham, MA, USA) connected to a QE-HF mass spectrometer (Thermo Fisher Scientific, Waltham, MA, USA). The applied protocol was adopted from [2]. In brief: 5 µl of the metabolite extract were injected in push partial loop mode using an overfill factor of 3, onto a Dionex IonPac AS11-HC column (2 mm × 250 mm, 4 μm particle size, Thermo Fisher Scientific, Waltham, MA, USA) equipped with a Dionex IonPac AG11-HC guard column (2 mm × 50 mm, 4 μm, Thermo Fisher Scientific, Waltham, MA, USA). The column temperature was held at 30°C, while the auto sampler was set to 6°C. The metabolites were separated using a potassium hydroxide gradient, which was generated by an eluent generator using a potassium hydroxide cartridge that was supplied with deionized water. The AEC flow rate was set to 380 µl/min, applying the following gradient. 0-8 min, 30-35 mM KOH; 8-12 min, 35-100 mM KOH; 12-15 min, 100 mM KOH, 15-15.1 min, 100-30 mM KOH. The column was re-equilibrated at 30 mM KOH for 3.9 min. The eluting metabolites were detected in negative ion mode using a MRM method on a Waters TQ triple quadrupole mass spectrometer (Waters). The MS settings for the analysis were: capillary voltage 2.75 kV, desolvation temperature 550°C, desolvation gas flow 800 l/h. The following MRM transitions were used for relative compound quantification of UDP and UDP-13C-Glc, respectively: m/z precursor mass (M-H^+^) 403/571, fragment mass (M-H^+^) m/z 159/323 using a cone voltage of 34/38 V and a collision energy of 26/24 V. For confirmation purpose of the analyzed compound an additional fragment mass (M-H^+^) m/z 111/96 using a cone voltage of 34/38 V and a collision energy of 20/34 was analyzed.

Data analysis and peak integration was performed using the TargetLynx Software (Waters). Both uridine and UDP was quantified using an independently generated calibration curve of the pure reference compound.

### Statistical Analyses

Depending on their genotype, littermate mice were allocated into experimental groups (P2Y6^fl/fl^ vs P2Y6^ΔNcr1^ or P2Y6^fl/fl^ vs P2Y6^ΔLysM^). For investigations of UDP and uridine levels, two independent cohorts of C57Bl6 mice were randomly assigned to HFD or CD feeding. The numbers of mice, representing the biological replicate, are included in the figure legends. Where multiple trials of the same experiment were conducted (technical replicates), such as for qPCR (duplicates), BMDM culture (duplicates/triplicates) and immunostainings (duplicates/triplicates), the results of these technical replicates were averaged and finally plotted as a single replicate. Statistical significance was tested using Graph Pad Prism 9.0.2 (GraphPad Software Inc., San Diego, CA, USA) and defined as significant, if p≤0.05. Statistical tests that were applied and resulting significance levels are indicated on figures and figure legends.

### Data Availability

Raw RNASeq data have been deposited in the NCBI Gene Expression Omnibus under accession code GSE175591.

## Acknowledgements

We thank all co-workers who supported this work at our institution, with special thanks to P. Scholl, N. Spenrath, C. Schäfer, K. Marohl, J. Noe and J. Frère for excellent technical assistance. We thank Dr. T. Wunderlich and Dr. C. Brandt for fruitful scientific discussions and Dr. H. Broenneke, Dr. U. Lichtenberg, Dr. K. Schoefisch and Dr. R. Braun for skillful administrative help.

## Abbreviations

µCT: Micro computer tomography
AgRP: Agouti-related protein
AKT: Protein kinase B
Apoa5: Apolipoprotein A-V
ATP: Adenosine triphosphate
BAT: Brown adipose tissue
BMDM: Bone marrow-derived macrophages
Ccl3: Chemokine (C-C motif) ligand 3
Ccl4: Chemokine (C-C motif) ligand 4
CD: Control Diet
CLS: Crown-like structures
cNK: conventional NK cells
Cxcl16: Chemokine (C-X-C motif) ligand 16
DAG: Diacylglycerides
DN: Double negative
ECAR: Extracellular acidification rate
OCR: Oxygen consumption rate
ELISA: Enzyme-linked immunosorbent assay
ERK: Extracellular signal-regulated kinases
FFA: Free fatty acids
GIR: Glucose infusion rate
GTT: Glucose tolerance test
HFD: High fat diet
HGP: Hepatic glucose production
HOMA-IR: Homeostasis model assessment- insulin resistance
HSL: Hormone-sensitive lipase
IFNγ: Interferon gamma
IL: Interleukin
ILC: Innate lymphoid cell
ITT: Insulin tolerance test
Lilra5: Leukocyte immunoglobulin like receptor A5
Lpl: Lipoprotein lipase
MAPK: Mitogen-activated protein kinase
NCD: Normal chow diet
NK cell: Natural killer cell
OCR: Oxygen consumption rate
OXPHOS: Oxidative phosphorylation
P2Y6R: P2Y purinoreceptor 6
PGAT: Perigonodal adipose tissue
pHSL: phosphorylated hormone-sensitive lipase
PI3K: Phosphoinositide 3-kinase
PKC: Protein kinase C
Plin2: Perilipin 2
RER: Respiratory exchange rate
Rora: RAR-related orphan receptor alpha
SEM: Standard error of the mean
SKM: Skeletal muscle
SVF: Stromal vascular fraction
TG: Triglycerides
TNFα: Tumor necrosis factor alpha
TPM: Transcripts per million
trNK: tissue-resident NK cells
UDP: Uridine diphosphate
uNK: uterine NK cells
Xcl1: Chemokine (C motif) ligand

